# Sex-dependent chronic neurological dysfunction following isoflurane anesthesia and surgery is associated with circulating extracellular vesicle-mediated neuroinflammatory signaling

**DOI:** 10.64898/2026.07.31.742052

**Authors:** Yun Li, Ruth Park, Balaji Krishnamachary, Hangnoh Lee, Zhuofan Lei, Hui Li, Junfang Wu

## Abstract

**Purpose:** It is well established that volatile anesthetics and surgery induce acute and subacute changes in the cellular and molecular landscape of the brain and peripheral circulation and can impair neurological function. However, the chronic neurological sequelae of isoflurane (Iso) anesthesia combined with surgical operation (OP), as well as the underlying mechanisms of postoperative neurological dysfunction, remain poorly understood.

**Methods:** Young adult male (M) and female (F) C57BL/6 mice underwent 4 h of 2% Iso plus laparotomy or sham treatment. At 12 weeks, olfactory and cognitive function were assessed by odor memory, buried food, Y-maze, and active avoidance tests. Olfactory bulbs (OB) and hippocampi (HI) were collected for RNAseq, while plasma extracellular vesicles (EVs) were isolated, characterized by NanoFCM, and profiled by Olink proteomics. Lastly, EVs were injected into the HI of naïve male mice, and cytokine/chemokine responses were measured 24 h later.

**Results:** Both sexes showed olfactory impairment after chronic Iso/OP, with greater deficits in females. Iso/OP mice, especially females, exhibited impaired odor recognition in the OM test and longer latencies to locate buried food. Female mice also showed greater hippocampal-dependent spatial working memory deficits in the Y-maze, with more arm returns and fewer alternations than Sham/F mice, whereas Iso/OP/M mice performed similarly to controls. Likewise, female, but not male, Iso/OP mice displayed impaired associative learning in the active avoidance test, evidenced by fewer avoided trials and more escape responses. These long-term behavioral abnormalities were accompanied by sex-divergent transcriptomic remodeling in the OB and HI, including altered synaptic, neurodevelopmental, extracellular matrix, stress-response, and chemotaxis-related pathways. Iso/OP reduced plasma EV particle numbers in both sexes and shifted EV size distributions, with prominent reductions in the 40-100 nm EV fraction. Proteomics revealed distinct sex-and condition-specific EV profiles, with several EV-associated proteins showing opposing sex-dependent expression patterns. Hippocampal injection of Iso/OP-derived EVs induced donor sex-dependent cytokine remodeling, confirming inflammatory bioactivity.

**Conclusions:** Four-hour isoflurane (Iso) exposure combined with laparotomy in young adult mice induces chronic, sex-dependent neurological deficits with distinct transcriptomic remodeling across brain subregions. Persistent alterations in circulating EV abundance and inflammatory cargo may drive chronic neuroinflammation and long-term brain dysfunction.

## 1 Introduction

Volatile anesthetics remain essential in clinical and experimental anesthesia [1, 2], yet increasing evidence suggests that anesthesia and surgery can produce persistent neurological effects [3–5]. Although isoflurane (Iso) has been replaced by sevoflurane in many operating rooms, it remains widely used in intensive care, resource-limited settings, and preclinical research [6–16]. Because general anesthesia is almost always accompanied by surgery and systemic inflammation, prolonged Iso exposure combined with laparotomy operation (OP) provides a clinically relevant model to investigate long-term neurological consequences. Although postoperative neurocognitive dysfunction has been studied mainly in older adults [17–20], prolonged anesthesia is also common in younger patients undergoing major surgery. Young adult models allow assessment of anesthesia-and surgery-induced neurological changes independent of aging.

Previous studies have shown that Iso/surgery induces acute and subacute cognitive and olfactory dysfunction accompanied by neuroinflammation and synaptic impairment [5, 20, 21]. Long-term neurological outcomes are clinically important because anesthesia and surgery may induce persistent changes beyond early postoperative recovery. Although clinical evidence for lasting neurological dysfunction remains mixed [22–24], preclinical models allow direct evaluation of long-term behavioral and molecular consequences. The olfactory bulb (OB) and hippocampus (HI), key regions for olfaction, memory, and learning, are particularly vulnerable to perioperative injury [25]. Consistent with our previous transcriptomic studies in aged mice, profiling these regions links persistent behavioral deficits to underlying molecular changes [21, 26].

Extracellular vesicles (EVs) have emerged as key mediators of intercellular communication and a potential link between peripheral inflammation and central nervous system (CNS) dysfunction [27]. EVs are membrane-bound nanoparticles that carry proteins, lipids, and nucleic acids reflective of their cells of origin and can cross the blood-brain barrier to deliver bioactive cargo to recipient cells [28, 29]. Circulating EVs contribute to neuroinflammatory signaling in neurological disorders [30–32], and their abundance and cargo are altered after anesthesia and surgery [26, 33–35], suggesting a role in postoperative neurological dysfunction. Whether they mediate persistent neuroinflammation after prolonged anesthesia and surgery remains unknown.

Biological sex is a recognized modifier of neurological outcomes after general anesthesia and surgery, but the direction of vulnerability remains inconsistent. Clinical studies have reported higher early postoperative delirium risk in male cardiac surgery and hip fracture patients [36, 37], whereas other analyses suggest greater risk in females [38]. Experimental studies show sex-dependent anesthetic effects: female 5xFAD mice developed cognitive deficits after sevoflurane and surgery, whereas neonatal male rats exhibited greater long-term impairment after Iso exposure [39, 40]. Sex also affects anesthetic sensitivity during Iso anesthesia in mice [41]. Whether prolonged anesthesia and surgery induce sex-dependent alterations in brain transcriptomes and circulating EV signaling that contribute to chronic neurological dysfunction has not been systematically investigated.

In the present study, we used a mouse model of prolonged Iso anesthesia combined with laparotomy to investigate long-term neurological outcomes in young adult male and female mice. We assessed long-term behavioral, transcriptomic, and circulating EV changes and evaluated the inflammatory bioactivity of plasma EVs in naïve recipient mice. We hypothesized that prolonged Iso/OP induces long-term, sex-dependent neurological dysfunction through altered EV-mediated neuroinflammatory signaling.

## 2 Materials and Methods

### 2.1 Animals and isoflurane/surgery (Iso/OP) model

Young adult (10-12-week-old) male (M) and female (F) C57BL/6 mice were obtained from Jackson Laboratories. All mice were housed in the same room of the animal care facility at the University of Maryland School of Medicine under a 12 h light/dark cycle, with ad libitum access to food and water. General anesthesia was induced and maintained with 2% Iso in 100% oxygen for 4 h as previously described [21, 26, 42]. Anesthesia depth was monitored through respiratory parameters (rate and depth) and reflex responses (palpebral and pedal). Core body temperature was maintained using a heated pad throughout the procedure. Following induction, mice were transferred to a surgical station with a nose cone for continuous Iso maintenance at 2% concentration. A midline laparotomy was performed by making a 1.5 cm incision from the xiphoid process to 0.5 cm above the pubic symphysis, sequentially penetrating the skin, abdominal musculature, and peritoneum. Following surgery, the incision site was treated with 0.25% bupivacaine in sterile saline, then closed in layers using 5-0 monofilament nylon sutures. The surgical procedure lasted approximately 10 to 15 min, after which mice were returned to the anesthesia chamber to complete the 4-h Iso exposure. During recovery, animals were placed on a temperature-regulated pad for 30-60 min to stabilize core temperature. Postoperative monitoring included intensive observation for 4 h post-Iso/OP and daily assessments thereafter to check on general well-being, incision site recovery and hydration.

### 2.2 Neurological function assessment

Following Iso/OP, a battery of neurological behavioral tests was conducted at the endpoint time of 12 weeks. The researcher conducting the behavioral experiments was blinded to group assignments until all behavior experiments were completed.

#### Odor memory (OM) test

To assess olfactory learning and memory, mice were individually housed overnight and tested using a previously established method [43, 44]. Briefly, a clean, dry, cotton-tipped wooden applicator (6 inches long) was inserted through a hole in the cage lid, allowing the mouse to familiarize itself with the applicator for 30 min. A fresh applicator was used for each subject. Cinnamon powder (McCormick, 100 ng/ml) was dissolved in water and stored in tightly sealed vials when not in use. During the first trial (T1), the mouse was exposed to the novel odor stimulus. The cotton tip of the applicator was dipped into the odor solution for 2 s before being inserted through the cage lid to a depth of approximately 2.5 cm. The total time the mouse spent sniffing the tip during the 5-minute trial (only when oriented toward the tip with its nose within 2 cm) was recorded. At the end of the trial, the applicator was removed. After a 60-min rest period, the second trial (T2) was conducted under the same conditions. The relative sniffing time ratio between trials, calculated as [T2/T1] ×100%, was used to assess memory retention. A reduced sniffing time during T2 indicated recognition and memory of the familiar odor.

#### Buried food (BF) test

To assess olfactory function, the BF test was performed to evaluate the ability of mice to detect and locate a familiar food item hidden beneath the bedding. This test measures the latency of an animal to find a buried, visually inaccessible food pellet using olfactory cues [44]. Briefly, mice were individually housed with ad libitum access to water and food-deprived for 24 h before testing to increase motivation for food seeking. The night before testing, a mini cookie was placed in each home cage for odor familiarization, and consumption was confirmed the following morning to verify palatability. On the test day, each mouse was placed in a clean test cage (46 cm length × 23.5 cm width × 20 cm height) containing 3 cm of fresh bedding and allowed to acclimate for 10 min before returning to its home cage. A uniformly sized mini cookie (∼1 g) was then buried 2-3 cm beneath the bedding in a randomly selected corner of the test cage. After the bedding surface was smoothed, the mouse was reintroduced into the test cage, and the latency to locate and begin consuming the buried cookie was recorded. The test was terminated if the mouse failed to locate the cookie within 15 min.

#### Open Field

The open-field test was used to evaluate spontaneous locomotor activity and anxiety-like behavior under dim-light conditions [21, 45]. At the beginning of the test, each mouse was placed individually in a corner of a 40 × 40 cm open-field arena, facing the wall, and allowed to explore freely for 5 min. Using the ANY-maze tracking system, the arena was divided into 10 × 10 cm grid squares. The peripheral 10-cm-wide area was designated as the outer zone, whereas the central 20 × 20 cm area was defined as the inner zone. Total distance travelled, average and maximum speed, and time spent in the inner and outer zones were recorded and analyzed using ANY-maze software (Stoelting Co., Wood Dale, IL, USA).

#### Rotarod Test

Motor coordination and locomotor function were assessed using an accelerating Rotarod apparatus (Harvard Apparatus), as described previously [46, 47]. Each mouse was placed on the stationary rod before the start of the trial. The Rotarod accelerated from 4 to 40 rpm over 90 s, with a maximum trial duration of 300 s. The latency to fall from the rotating rod was recorded for each trial. Each mouse completed five trials, and the mean latency to fall was calculated and used for between-group comparisons.

#### Grip Strength

Grip strength was assessed to evaluate neuromuscular function, particularly forelimb muscle strength, using a digital grip strength meter (Ugo Basile) according to our established protocols [46, 47]. Briefly, the forelimb GS of both paws was assessed by placing the mouse on a mesh wire grid connected to the device’s force transducer. Once the mouse secured a firm grip, it was gently held by the tail and slowly pulled away from the grid. The maximum force exerted on the mesh wire grid was recorded, with each mouse undergoing an average of 10 daily trials. Final GS values were normalized to body weight for comparison between groups.

#### Y maze (YM) test

The Y-maze test was performed to assess hippocampus-dependent spatial working memory in mice, as described previously [47–49]. The Y-maze apparatus (Stoelting Co.) consisted of three arms of identical length and width, designated A, B, and C. During the test, one arm was randomly selected as the starting position, and each mouse was placed in the maze and allowed to explore freely for 5 min. Arm entries were recorded, and a spontaneous alternation was defined as consecutive entries into three different arms. The percentage of spontaneous alternation was calculated using the following equation: total alternations × 100/ (total arm entries – 2). Mice with alternation percentages significantly above 50%, the chance level for entering a novel arm, were considered to have intact spatial working memory.

#### Active Avoidance test

Associative learning was evaluated using a revised version of the two-way active avoidance task in a Gemini shuttle box apparatus, as described by Macheda et al [50]. Mice were moved to the testing room and allowed to acclimate for at least 30 min before testing. The shuttle box consisted of two compartments separated by an open guillotine door. At the beginning of each session, each mouse was placed into one compartment and allowed to habituate freely for 5 min. Each daily session consisted of 30 trials. During each trial, a conditioned stimulus (house light, 10 s) was presented, followed by an unconditioned stimulus (0.2 mA foot shock, 2 s) unless the mouse crossed to the opposite compartment during the CS period. The intertrial interval was randomly varied between 25 and 35 s, with an average interval of 30 s. Crossings during the CS period were scored as avoidance responses, whereas crossings after shock onset were scored as escape responses. Trials in which mice failed to cross during the shock period were scored as escape failures. Avoidance responses, escape responses, and escape failures were quantified to assess associative learning performance. The chamber floor and walls were cleaned between mice to maintain consistent testing conditions

### 2.3 RNA extraction and bulk RNA sequencing (RNAseq)

Following neurological functional assessment, mice were euthanized, blood samples were collected, and the mice were perfused with 50 mL of ice-cold saline. Total RNA was isolated from dissected OB and HI tissues of mice using the RNAeasy mini kit (Cat# 74106, Qiagen). The RNA samples were sent to Novogene (Sacramento, CA) for mRNA library preparation (poly A enrichment) and paired-end sequencing (150 bp) on a NovaSeq 6000 platform (Illumina, CA). We aligned RNAseq reads to the mouse reference genome (GRCm39) using STAR (version 2.7.5) [51] with GENCODE gene annotation (version M33). Gene and isoform expression levels were quantified in transcripts per million (TPM) using RSEM (version 1.3.3) [52]. Differential expression gene (DEG) analysis was performed with DESeq2 (version 1.42.1) [53] and false discovery rates (FDR) were calculated using the Benjamini-Hochberg method. For partial least squares discriminant analysis (PLS-DA), the package Mixomics was used. Genes with an FDR of less than 0.05 were differentially expressed and used for downstream pathway enrichment analysis. Gene ontology (GO) analysis was performed with cluster Profiler 4.10.1 using DESeq2 output [54].

### 2.4 Plasma EV isolation

Blood was collected into precoated EDTA tubes (Cat #365974, BD Biosciences) through terminal cardiac puncture under anesthesia. EVs were isolated using differential ultracentrifugation, with minor modifications as described in our previous publications [26, 31, 48, 55]. Briefly, blood samples were immediately centrifuged at 800 *xg* for 10 min, followed by 2500 *x g* for 10 min, to generate platelet free plasma. To separate larger EVs, equal volumes of plasma from each animal were centrifuged at 12,000 *xg* for 30 min at 4 °C. The supernatant was then subjected to ultracentrifugation at 110,000 *xg* for 2 h at 4 °C to isolate small EVs. The pellet was washed with phosphate buffered saline (PBS) and re-centrifuged under the same conditions. The final EV pellet was resuspended in an equal volume of PBS.

### 2.5 Nano flow cytometry

Particle concentration and size of EVs were measured using a Flow NanoAnalyzer (NanoFCM)[26]. The instrument was calibrated using QC and size standard beads. EV samples were diluted in sterile, molecular-grade water, and the events were recorded for 1 minute. Particle concentration and size were calculated using the calibration curves[56]. To detect canonical tetraspanin markers on EVs, 5μL of EV sample was incubated with APC-conjugated antibodies against CD63 (1:10, Cat#143905, BioLegend), CD9 (1:10, Cat#124812, BioLegend), or CD81 (1:10, Cat#104909, BioLegend). For CD9 and CD63, 3 μL of diluted antibody was used, whereas 1 μL was used for CD81. Samples were incubated for 30 min at 37 °C at 60 RPM, followed by dilution in molecular-grade water and acquisition on a NanoFCM instrument. Molecular-grade water alone and antibody-only preparations were used as blank and staining controls, respectively. A total of 2,000-12,000 events were recorded per sample. Particle concentration was calculated from flow rate and side-scatter intensity using NanoFCM Profession software (v3.0).

### 2.6 Plasma EVs proteomics and data analysis

Plasma EVs were lysed in radioimmunoprecipitation assay buffer for protein quantification. Equal amounts of EV protein (0.5 μg/μL) were loaded in randomized order into a 96-well plate and submitted to Psomagen (Rockville, MD, USA) for proteomic analysis as described in previous publication [55]. EV protein cargo was profiled using the Olink Mouse Exploratory Panel. Protein detection was performed using Olink’s proximity extension assay, a multiplex immunoassay in which pairs of oligonucleotide-labeled antibodies bind to their target proteins. Upon dual binding, the oligonucleotides hybridize and form a DNA template that is subsequently quantified by real-time PCR. Samples were randomly plated, and assay performance was monitored using internal quality controls. Protein abundance was reported as normalized protein expression (NPX) values on a log2 scale, with higher NPX values indicating higher relative protein levels. Following the manufacturer’s guidelines, samples that did not meet the technical specifications of the assay were excluded from further analysis. Pairwise comparisons were performed using the limma framework in R, fitting linear models to NPX values followed by empirical Bayes shrinkage to obtain moderated statistics for differential protein expression [57, 58].

### 2.7 Processing of donor plasma EV, intra-hippocampal injection, and cytokine array

The donor plasma EV samples were pooled within each group and stored at −80 °C until required for injections. For the intra-hippocampal EV injection experiments, young adult C57BL6 male mice were anesthetized by Iso and placed in a stereotaxic setup. Artificial eye ointment was applied to protect the eye, and Bupivacaine was injected subcutaneously as local analgesia. Following a midline scalp incision, two small holes were drilled on the skull above the left hippocampus for each mouse (dorsal hippocampus: AP −2.0 mm, ML −1.5 mm, DV 2.0∼1.8 mm; Ventral hippocampus: AP −3.0 mm, ML −2.8 mm, DV 3.0∼2.8 mm). Each mouse received 0.5 μl EV injection per site through a 5 μl syringe (Hamilton™ Neuros™ 700 Series). The injection volume and flow rate were controlled as 0.5 μl at 0.1 μl/min. The needle was moved up 0.1 mm per 0.25 μl during the injection and was kept in place for 3 additional min and then slowly withdrawn. The mice were sutured with Bupivacaine, reapplied and then returned to their home cages for recovery. All animals were sacrificed for hippocampus tissue collection 24 h later. For cytokine profiling, hippocampal tissues from each group were lysed using the lysis buffer provided in the Mouse Cytokine Array C3-(AAM-CYT-3-8, Ray Biotech). The samples were pooled to minimize inter-individual variability. A total of 500 µg of protein per group was used for the cytokine array. Samples were incubated with the membrane overnight at 4 °C according to the manufacturer’s protocol. Each cytokine/chemokine was spotted in duplicate on the array membrane. The experiment was performed in two independent runs using separate membranes. Membrane images were analyzed using Bio-Rad Image Lab software. Cytokine array spot intensities were first normalized to the positive control spots on each membrane. Values were then normalized to the male sham EV-injected group as the reference control. For each analyte, duplicate spots from two independent membranes were combined, yielding four values per cytokine/chemokine for downstream analysis and heatmap generation. For visualization, normalized values were converted to row-scale z-scores for each analyte.

### 2.8 Statistical Analysis

All quantitative data are displayed as individual data points in column graphs, presented as mean ± SEM. Statistical analyses were conducted using GraphPad Prism (version 9.5.0 for Windows, GraphPad Software; RRID: SCR_002798) for all tests. The Shapiro-Wilk test was used to assess data distribution normality. For multiple group comparisons, one-way ANOVA or repeated-measures two-way ANOVA were performed, followed by Tukey’s or Newman-Keuls post hoc tests for parametric data (when normality and equal variance assumptions were met). Detailed statistical analyses for each assay are provided in the figure legends, with a significance threshold set at p ≤ 0.05. repeated-measures two-way ANOVA with Tukey’s multiple-comparisons test.

## 3 Results

### 3.1 Extended Iso/OP induces chronic sex-dependent post-operative neurobehavioral deficits

To assess chronic neurological dysfunction, young adult male and female C57BL/6 mice underwent abdominal surgery followed by 4 h of Iso anesthesia. A battery of behavioral tests was performed 12 weeks after Iso/OP to evaluate long-term neurobehavioral outcomes. Odor memory was first assessed using the odor recognition test. Mice were initially exposed to a cotton swab soaked in a cinnamon solution, a mildly aversive odor, and were re-exposed to the same odor at least 1 h later. Mice that successfully remembered the odor displayed reduced investigation during the second trial. Baseline odor memory was comparable between male and female sham mice, with no significant sex differences detected. At 12 weeks after Iso/OP, both male and female mice exhibited impaired odor memory compared with their respective sham controls; however, no significant sex difference was observed between the Iso/OP groups (Fig. 1A). We next evaluated odor sensitivity using the buried food test. Consistent with the odor memory results, sham male and female mice performed similarly, whereas Iso/OP impaired odor sensitivity in both sexes (Fig. 1B). Notably, Iso/OP/F mice required significantly more time to locate the buried food pellet than Iso/OP/M mice, indicating greater olfactory dysfunction in females (Fig. 1B). These findings suggest that Iso/OP induces chronic olfactory deficits lasting at least 12 weeks after surgery, with female mice exhibiting greater impairment in odor sensitivity. To assess long-term cognitive function, hippocampus-dependent spatial working memory was evaluated using the Y-maze. Iso/OP/F mice displayed significantly reduced spontaneous alternation compared with Sham/F controls, whereas no significant impairment was detected in male mice (Fig. 1C-D). Similarly, active avoidance testing, which measures associative learning and memory, revealed a significant reduction in avoidance performance in Iso/OP/F mice but not in Iso/OP/M mice (Fig. 1E-F). Collectively, these findings demonstrate that extended Iso/OP induces chronic neurobehavioral deficits in young adult mice. Although both sexes exhibited long-term olfactory impairments, female mice were more susceptible to Iso/OP-induced deficits, displaying greater impairment in odor sensitivity, spatial working memory, and associative learning.

**Fig. 1.**
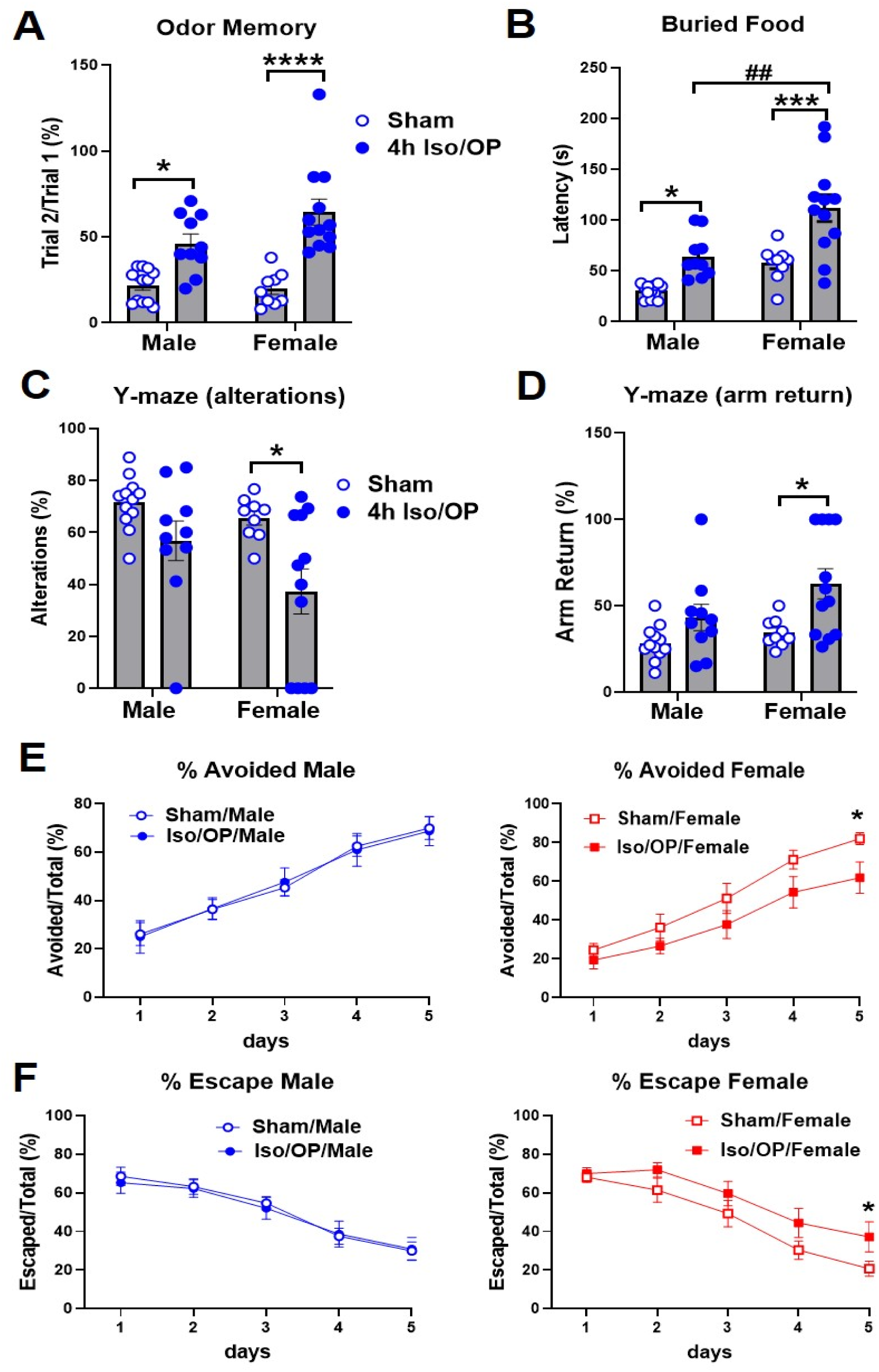
Four-hour isoflurane exposure with laparotomy induces chronic sex-dependent neurobehavioral deficits in young adult C57BL/6 mice. Young adult male and female mice (8-11 weeks old) were subjected to laparotomy under 4 h isoflurane exposure (Iso/OP), followed by behavioral analyses 3 months after exposure. **A-B** Olfactory function was assessed using the odor memory test **(A)** and buried food test **(B)**. **C-D** Spatial working memory was assessed using the Y-maze test, including spontaneous alternations **(C)** and arm returns **(D)**. **E-F** Active avoidance performance was assessed across 5 training days, including percentage avoided trials **(E)** and percentage escape trials **(F)**, shown separately for male and female mice. n = 9-12 mice/group. *p < 0.05, ***p < 0.001, and ****p < 0.0001 vs. sham controls; #p < 0.05 and ##p < 0.01 vs. Iso/OP male mice, as indicated. Data were analyzed by Student’s t-test, two-way ANOVA, or repeated-measures two-way ANOVA with Tukey’s multiple-comparisons test, as appropriate.

In addition, general locomotor activity and neuromuscular function were assessed (Supplemental Fig. S1). In the open field test, Sham/F mice traveled a greater total distance than Sham/M mice, indicating a baseline sex difference. However, Iso/OP did not affect any open field parameters (total distance, maximum speed, mean speed, or immobility time) in either sex. Likewise, rotarod performance and grip strength showed no sex-or Iso/OP-dependent differences. As expected, female mice weighed significantly less than males, but body weight was unaffected by Iso/OP in either sex. Together, these findings indicate that Iso/OP did not alter locomotor activity, neuromuscular function, or body weight under the conditions tested, while expected baseline sex differences were preserved.

### 3.2 Olfactory bulbs and hippocampus undergo sex-divergent transcriptomic remodeling after chronic Iso/OP

Because the long-term neurological impairments were primarily localized to the OB and HI, we next examined molecular changes in these regions using bulk RNAseq at 12 weeks after Iso/OP cessation. In the OB, PLS-DA of normalized gene counts showed clear separation among the four groups (Fig. 2A). Differential expression analysis (FDR < 0.05) identified only modest transcriptional differences between Iso/OP/F and Iso/OP/M mice, with the most significant DEGs primarily representing sex chromosome-linked genes, including *Kdm5d*, *Ddx3y*, *Uty*, and *Xist* (Fig. 2B). GO biological process analysis revealed significant enrichment only among downregulated DEGs (Fig. 2C). The top enriched pathways involved small GTPase-mediated signal transduction, synapse organization, and synaptic signaling, with representative genes shown in the accompanying heatmaps and log2 fold-change plots (Fig. 2D-E). Reactome analysis further identified cellular stress, the RHO GTPase cycle, and axon guidance as the most significantly regulated pathways (Fig. 2F), while WikiPathways highlighted MAPK signaling, mechanisms of pluripotency, and EGFR1 signaling as the principal pathways exhibiting sex-dependent regulation following Iso/OP (Fig. 2G).

**Fig. 2.**
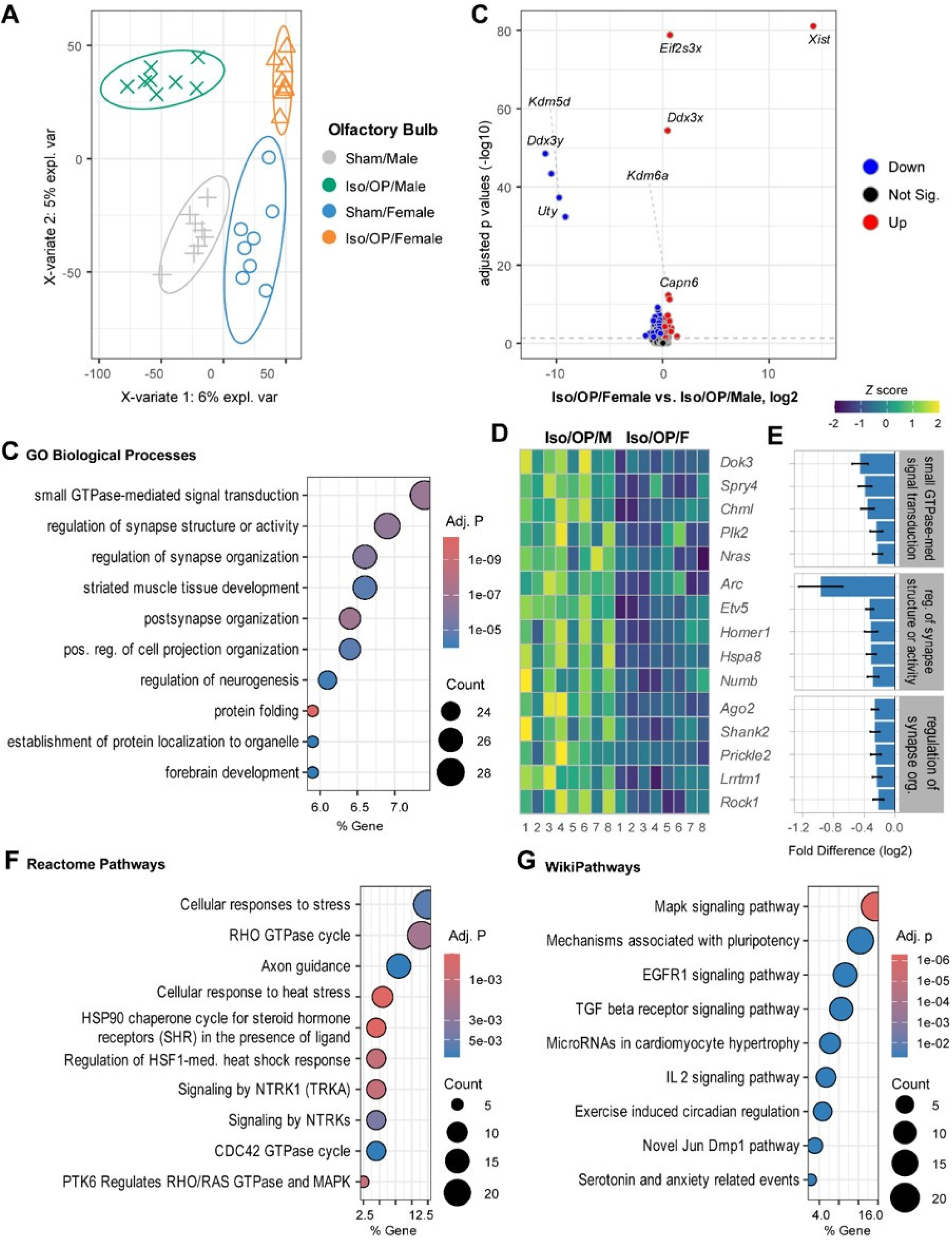
Four-hour isoflurane exposure with laparotomy induces sex-divergent transcriptomic remodeling in the olfactory bulb (OB) of young adult mice. Young adult male and female C57BL/6 mice (8-11 weeks old) were subjected to laparotomy under 4 h isoflurane exposure (Iso/OP), followed by euthanasia and OB dissection 3 months after exposure. **A** PLS-DA of normalized OB transcriptomic counts from sham and Iso/OP male and female mice. **B** Volcano plot of differentially expressed genes (DEGs) in Iso/OP/Female vs. Iso/OP/Male mice. **C-E** GO biological process enrichment of DEGs downregulated in Iso/OP/Female relative to Iso/OP/Male mice **(C)**, with heatmap visualization of representative signature genes from selected enriched pathways **(D)** and fold-difference plots for representative pathway genes **(E)**. **F-G** Reactome **(F)** and WikiPathways **(G)** enrichment analysis of downregulated DEGs in Iso/OP/Female relative to Iso/OP/Male mice. Dot size represents gene count, color indicates adjusted p value, and the x-axis shows gene ratio. RNAseq data were analyzed with DESeq2; n=8 mice/group.

Because Iso/OP caused persistent olfactory deficits that were more pronounced in female mice, we next examined baseline sex differences and sex-specific transcriptomic remodeling in the OB at 12 weeks after exposure. Comparisons were performed between Sham/F and Sham/M, Iso/OP/M and Sham/M, and Iso/OP/F and Sham/F groups. Baseline analysis revealed clear sex-associated differences in OB gene expression, with prominent differential expression of sex chromosome-linked genes, including *Xist*, *Ddx3y*, *Eif2s3x*, *Kdm5d*, and *Uba1y* (Supplemental Fig. S2A). GO analysis identified enrichment of pathways related to responses to external stimuli, vitamin metabolism, and terpenoid metabolism, while heatmap analysis further demonstrated distinct baseline transcriptional profiles between sexes (Supplemental Fig. S2B-E). In female mice, Iso/OP induced relatively few DEGs but prominently altered pathways involved in synaptic vesicle-mediated transport, vesicle organization and fusion, exocytosis, neurotransmitter release, and membrane fusion, together with enrichment of iron transport, estrogen receptor signaling, and purine metabolism (Supplemental Fig. S2F-H). Consistently, genes involved in synaptic vesicle function and neurotransmitter release, including *Snap25*, *Stx1b*, *Atp6v0e2*, *Atp6v1g2*, *Cplx1*, and *Doc2g*, were differentially expressed (Supplemental Fig. S2I-J). In contrast, male mice exhibited a broader transcriptional response to Iso/OP, with a larger number of DEGs and enrichment of pathways associated with peptide and hormone transport, PI3K/PKB signaling, cell migration, protein localization, and chemotaxis (Supplemental Fig. S2K-M). Representative altered genes included *Per2*, *Crhbp*, *Irs2*, *Vgf*, *Slc25a39*, *Dynll1*, and *Camk2n1* (Supplemental Fig. S2N-O). Together, these findings demonstrate baseline sex-specific transcriptional differences in the OB and distinct molecular responses to Iso/OP. Whereas female mice showed selective remodeling of synaptic vesicle and neurotransmitter transport pathways, male mice exhibited broader changes involving transport, PI3K/PKB signaling, and cell migration. These sex-dependent molecular differences may contribute to the greater olfactory impairment observed in female mice.

We next examined HI transcriptomic changes to determine whether the long-term cognitive deficits were accompanied by molecular remodeling in this key learning-and memory-related brain region. PLS-DA revealed clear separation between sham and Iso/OP groups, with distinct clustering of Iso/OP/F and Iso/OP/M samples (Fig. 3A). Direct comparison of Iso/OP/F and Iso/OP/M mice identified pronounced sex-dependent differences in hippocampal gene expression, including expected sex chromosome-linked transcripts (e.g., *Xist*, *Ddx3y*, *Kdm5d*, and *Uty*) (Fig. 3B). GO analysis showed that genes upregulated in Iso/OP/F versus Iso/OP/M mice were enriched in pathways related to metal ion response, glycerolipid metabolism, oxidative stress, learning, microtubule dynamics, phospholipid catabolism, extracellular matrix organization, and hyaluronan biosynthesis (Fig. 3C). Representative genes from these pathways, including *Aqp1*, *Mt2*, *Enpp2*, *Pdgfb*, and *Rgs14*, showed higher expression in the female hippocampus (Fig. 3D-E). In contrast, genes downregulated in Iso/OP/F relative to Iso/OP/M mice were enriched in protein folding, nucleocytoplasmic transport, neurogenesis, protein localization, cell projection organization, mRNA metabolism, forebrain development, and endoplasmic reticulum stress pathways (Fig. 3F), with representative male-enriched genes including *Hspa1b*, *Hspb1*, *Hspa5*, *Dnajb1*, *Pdia4*, *Xbp1*, *Drd2*, *Sox11*, and *Btg2* (Fig. 3G-H). Together, these findings demonstrate that Iso/OP induces persistent, sex-dependent hippocampal transcriptomic remodeling. Female mice preferentially upregulated pathways associated with oxidative stress, lipid metabolism, extracellular matrix remodeling, and learning-related processes, whereas male mice showed greater enrichment of protein-folding, nucleocytoplasmic transport, neurodevelopmental, and endoplasmic reticulum stress pathways. These molecular differences parallel the sex-specific behavioral impairments observed in spatial working memory and associative learning.

**Fig. 3.**
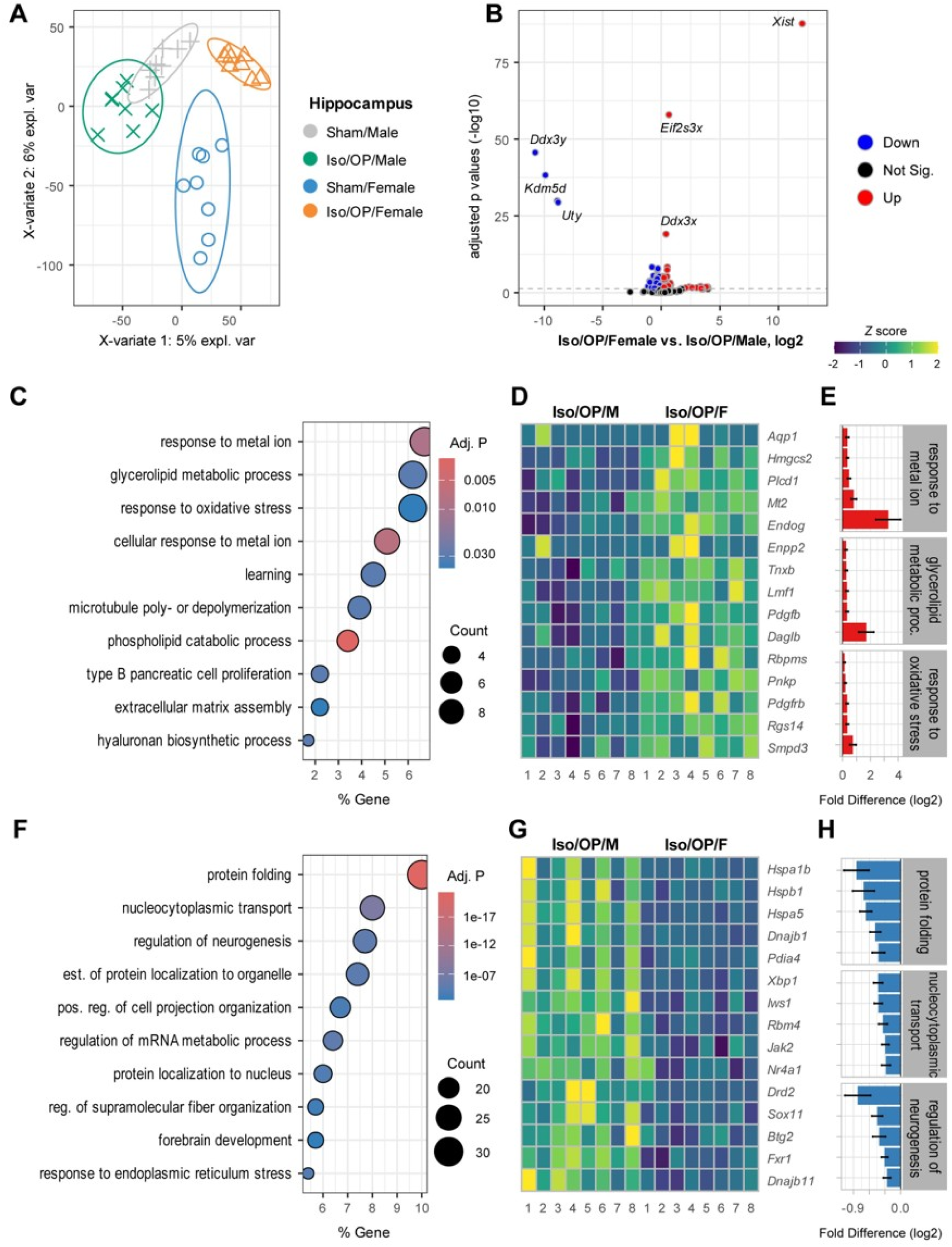
Four-hour isoflurane exposure with laparotomy induces sex-divergent transcriptomic remodeling in the hippocampus (HI) of young adult mice. **A** PLS-DA of normalized HI transcriptomic counts from sham and Iso/OP male and female mice. **B** Volcano plot of DEGs in Iso/OP/Female vs. Iso/OP/Male mice. **C-E** GO biological process enrichment of DEGs upregulated in Iso/OP/Female relative to Iso/OP/Male mice **(C)**, with heatmap visualization of representative signature genes from selected enriched pathways **(D)** and fold-difference plots for representative pathway genes **(E)**. **F-H** GO biological process enrichment of DEGs downregulated in Iso/OP/Female relative to Iso/OP/Male mice **(F)**, with heatmap visualization of representative signature genes from selected enriched pathways **(G)** and fold-difference plots for representative pathway genes **(H)**. Dot size represents gene count, color indicates adjusted p value, and the x-axis shows gene ratio. RNAseq data were analyzed with DESeq2; n=8 mice/group.

In the HI, comparison of sham female and sham male mice revealed clear baseline sex-dependent transcriptional differences, largely driven by expected sex chromosome-linked transcripts (Supplemental Fig. S3A). GO analysis showed enrichment of pathways related to body fluid regulation, glomerular filtration, astrocyte development, and epithelial/glial cell migration (Supplemental Fig. S3B), while heatmap analysis highlighted distinct baseline expression of genes involved in water transport, extracellular matrix organization, and structural remodeling (Supplemental Fig. S3C). In female mice, Iso/OP induced modest but detectable transcriptomic remodeling relative to sham controls (Supplemental Fig. S3D). Upregulated DEGs were enriched in developmental, gliogenic, dendritic spine, tissue remodeling, and axon guidance pathways (Supplemental Fig. S3E), whereas downregulated DEGs were associated with chemotaxis-and immune-related processes (Supplemental Fig. S3F). Representative genes, including *Fgfr3*, *Shank1*, *Cdk5r2*, *Dlgap4*, *Cst3*, *Plec*, *Shank3*, and *Scg2*, showed altered expression in Iso/OP females (Supplemental Fig. S3G-H). In contrast, Iso/OP produced minimal hippocampal transcriptional changes in male mice, with Top1 as the only significantly upregulated gene (Supplemental Fig. S3I). Accordingly, GO enrichment was limited and driven by a small number of genes, highlighting pathways related to embryonic cleavage, DNA conformation, rRNA transcription, circadian regulation, and responses to cAMP or organophosphorus compounds (Supplemental Fig. S3J). Heatmap analysis confirmed increased Top1 expression in Iso/OP males (Supplemental Fig. S3K). Together, these analyses demonstrate pronounced baseline sex differences in the hippocampal transcriptome and distinct sex-specific responses to Iso/OP. Female mice exhibited broader remodeling involving neurodevelopmental, synaptic, extracellular matrix, and immune pathways, whereas male responses were comparatively limited, supporting the primary findings of sex-dependent long-term hippocampal molecular remodeling after Iso/OP.

### 3.3 Chronic Iso/OP disrupts circulating EV profiles and protein cargo

To determine whether Iso/OP exposure and biological sex alter circulating EVs during the chronic post-surgical phase, plasma EVs were prepared 12 weeks after Iso/OP or sham exposure. NanoFCM analysis revealed significantly reduced plasma EV concentrations in both male and female Iso/OP mice compared with their respective sham controls (Fig. 4A). Two-way ANOVA further demonstrated that plasma EV concentrations were lower in Sham/F mice than in Sham/M mice and were further reduced in Iso/OP/F mice relative to Iso/OP/M mice (Fig. 4B). Size-distribution analysis showed that circulating EVs were predominantly enriched within the small EV range, with peak abundance between 40 and 100 nm (Fig. 4C). Across both sexes, Iso/OP reduced particle abundance around this major distribution peak compared with sham controls, with the greatest reduction observed in Iso/OP/M mice. Binned size analysis confirmed that the observed group differences were driven primarily by the 40-100 nm fraction, whereas larger particle populations were less abundant and exhibited greater variability (Fig. 4D). Collectively, these findings demonstrate that Iso/OP induces chronic, sex-dependent alterations in circulating EV abundance and size distribution, particularly within the small EV-enriched population.

**Fig. 4.**
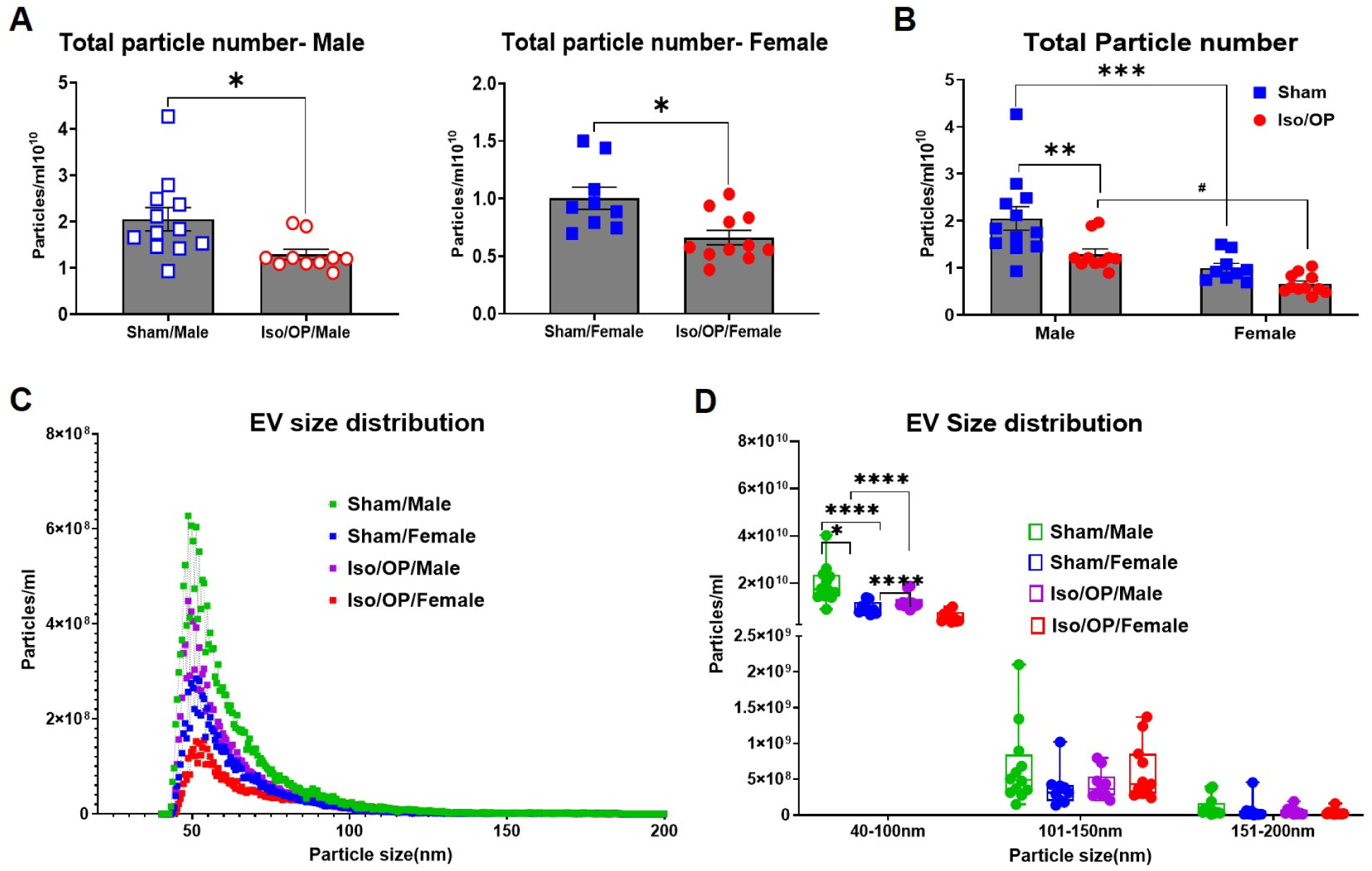
Four-hour isoflurane exposure with laparotomy alters circulating extracellular vesicle (EV) particle concentration and size distribution in young adult mice at 3 months after exposure. Plasma EV particle concentration and size distribution were analyzed using NanoFCM. **A** Total particle number in male and female mice analyzed within each sex. **B** Total particle number across sex and treatment groups. **C** EV particle size distribution curves across experimental groups. **D** Quantification of EV particle numbers by size range (40-100 nm, 101-150 nm, and 151-200 nm). n=9-12 mice/group. *p < 0.05, **p < 0.01, ***p < 0.001, and ****p < 0.0001 vs. indicated groups; #p < 0.05 vs. indicated group. Data were analyzed by Student’s t-test or two-way ANOVA with Tukey’s multiple-comparisons test, as appropriate.

Furthermore, EV-associated proteins were profiled using the Olink Mouse Exploratory panel 12 weeks after exposure. PLS-DA revealed clear separation among the four groups, indicating that both biological sex and Iso/OP exposure contributed to variation in EV protein composition (Fig. 5A). Notably, Iso/OP/M and Iso/OP/F samples separated along distinct axes, suggesting divergent EV protein signatures following Iso/OP. Differential expression analysis identified multiple EV-associated proteins that differed significantly between Iso/OP/M and Iso/OP/F mice (Fig. 5B). Among these, lipoprotein lipase (LPL) was markedly reduced in Iso/OP/F mice relative to Iso/OP/M mice and met both statistical significance and fold-change thresholds. Additional proteins, including CCL2, CCL20, CXCL1, PRDX5, FST, FLI1, WISP1, SNAP29, ADAM23, and PDGFB, also exhibited sex-or condition-dependent differences in EV abundance. Heatmap visualization of group-mean z-scores demonstrated coordinated remodeling of EV protein cargo across sham and Iso/OP groups, with distinct protein clusters enriched in male or female Iso/OP mice (Fig. 5C). Representative analyte plots further illustrated sex-dependent abundance patterns for selected EV-associated proteins (Fig. 5D). Several inflammatory and signaling molecules, including CCL2, CCL20, CXCL1, and WISP1, displayed differential abundance between male and female Iso/OP mice, whereas LPL, FST, PRDX5, and ADAM23 highlighted broader alterations related to lipid metabolism, oxidative stress, and neurodevelopmental processes. Together, these findings demonstrate that Iso/OP induces sustained remodeling of circulating EV protein cargo, generating distinct sex-dependent molecular signatures that may reflect divergent systemic and neuroimmune responses to anesthetic and surgical stress.

**Fig. 5.**
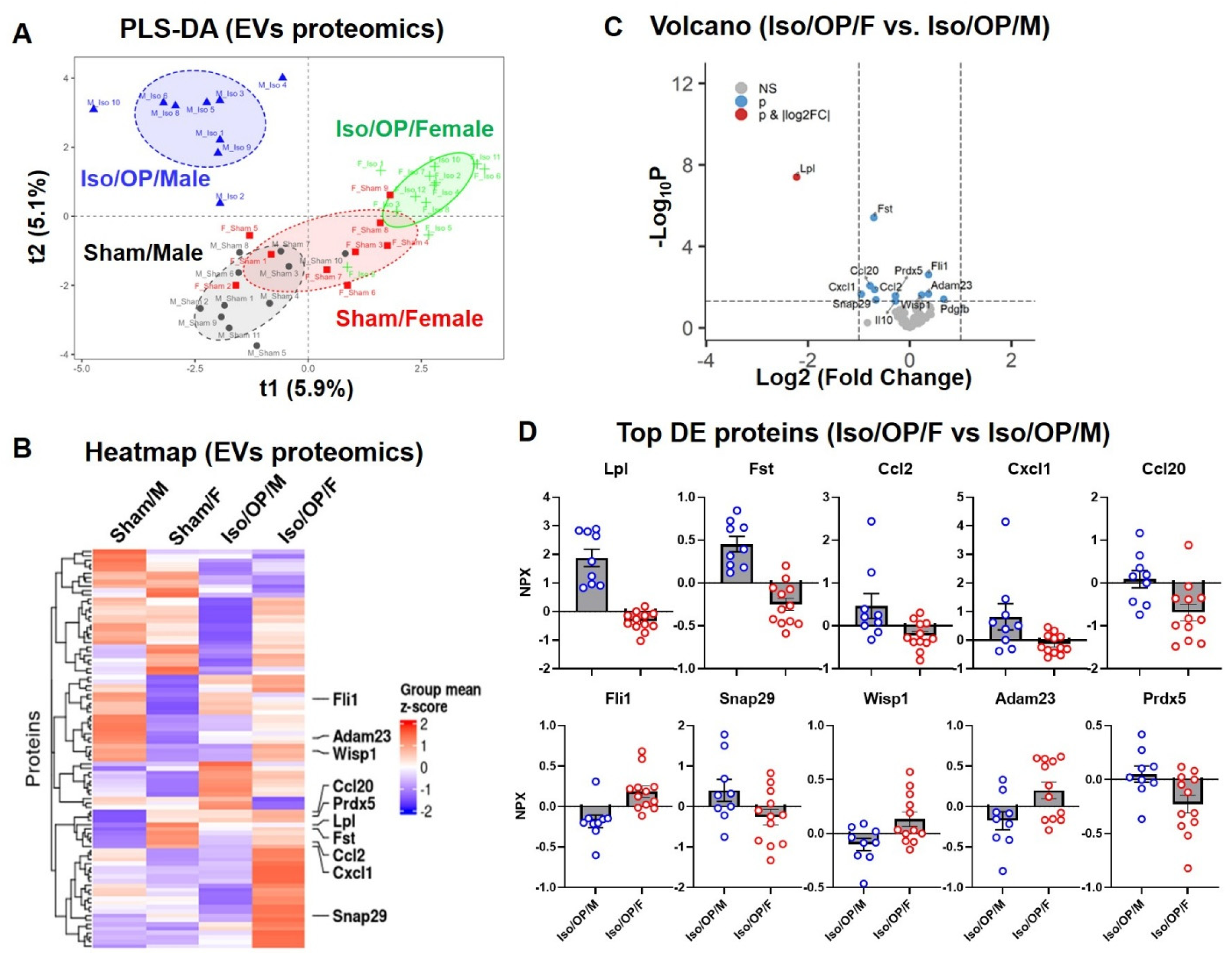
Circulating EV protein cargo is altered 3 months after 4 h isoflurane exposure with laparotomy in young adult mice. Plasma EV protein cargo was profiled using the Olink Mouse Exploratory panel. **A** PLS-DA of EV proteomic profiles from sham and Iso/OP male and female mice. **B** Group-mean z-score heatmap of EV protein abundance across experimental groups. **C** Volcano plot of differentially represented EV proteins in Iso/OP/Female vs. Iso/OP/Male mice. **D** Individual abundance plots of representative differentially represented EV proteins between Iso/OP/Female and Iso/OP/Male mice. n=9-12 mice/group.

To further investigate whether this sex-dependent remodeling preferentially involved inflammatory signaling, we next focused on cytokine-, chemokine-, and inflammation-related proteins identified in the EV Olink dataset. PLS-DA based on this subset of inflammatory EV-associated proteins demonstrated clear separation between Iso/OP/M and Iso/OP/F samples, indicating distinct inflammatory EV signatures following Iso/OP (Fig. 6A). Heatmap analysis likewise revealed coordinated, sex-dependent differences in inflammatory EV protein cargo across the groups (Fig. 6B). Several inflammatory mediators, including CXCL1, CCL2, IL-6, and CCL3, exhibited higher relative abundance in EVs from Iso/OP/M mice, whereas other immune-related proteins showed preferential enrichment in Iso/OP/F mice or sham-derived EVs. These findings suggest that Iso/OP does not globally increase or suppress EV-associated inflammatory proteins but instead selectively reshapes the inflammatory cargo of circulating EVs in a sex-dependent manner. Representative analyte plots further confirmed sex-and condition-dependent differences in selected inflammatory EV-associated proteins (Fig. 6C). Collectively, these data demonstrate that Iso/OP produces persistent remodeling of the inflammatory protein landscape carried by circulating EVs. These observations provided the rationale for subsequently testing whether plasma EVs isolated from Iso/OP mice possess functional inflammatory bioactivity capable of modulating inflammatory responses in brain tissue.

**Fig. 6.**
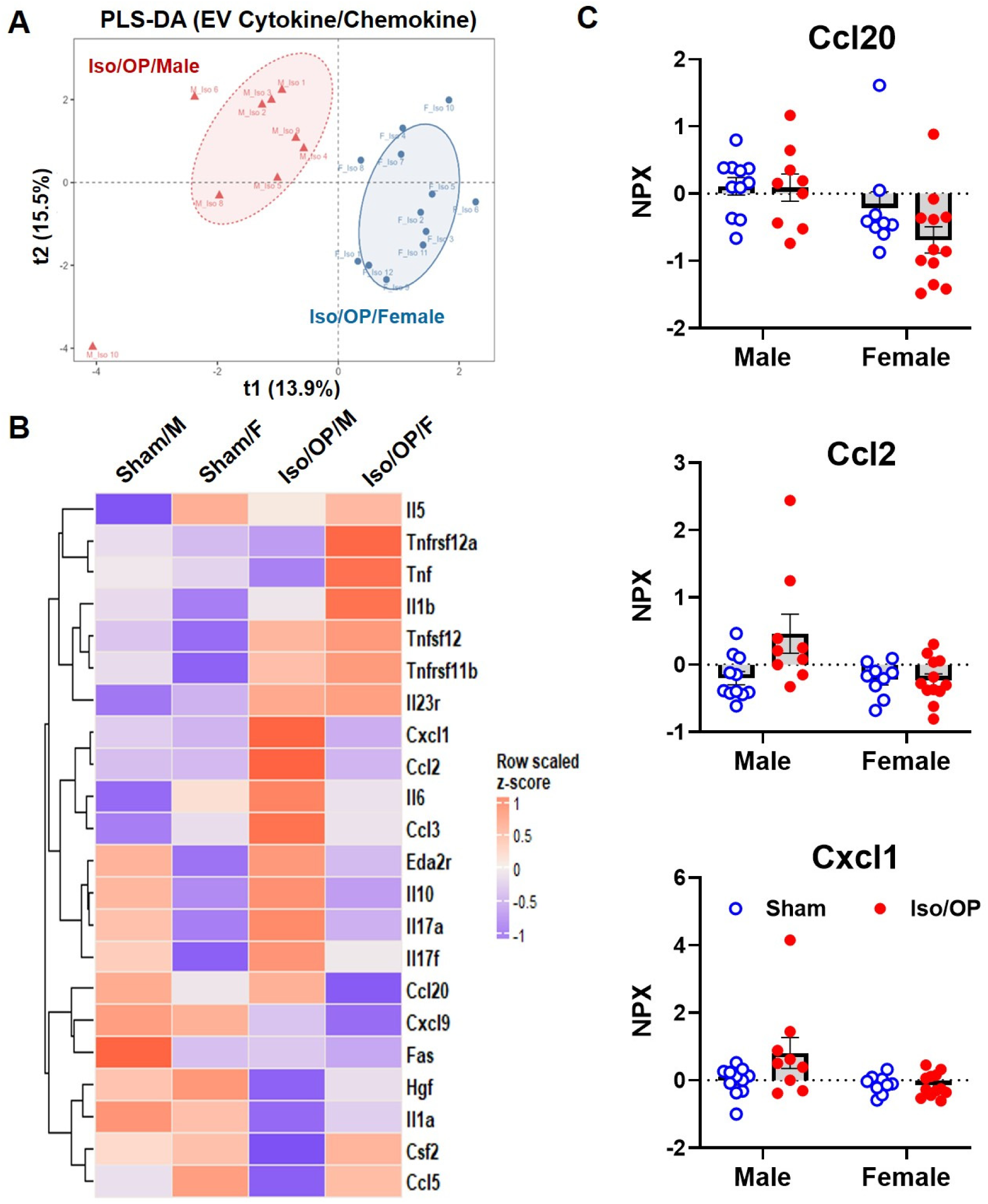
Iso/OP induces sex-divergent circulating EV cytokine and chemokine cargo profiles at 3 months after exposure. Circulating EVs were isolated from plasma collected 12 weeks after sham or Iso/OP exposure, and EV-associated inflammatory proteins were profiled using the Olink Mouse Exploratory panel. **A** PLS-DA score plot showing separation of Iso/OP male and Iso/OP female mice based on EV cytokine/chemokine cargo. Each point represents an individual animal; ellipses indicate group clustering. **B** Heatmap showing row-scaled z-scores of selected cytokine-, chemokine-, and inflammation-related EV cargo proteins across Sham/Male, Sham/Female, Iso/OP/Male, and Iso/OP/Female groups. **C** Representative EV-associated inflammatory proteins showing sex-and Iso/OP-associated abundance patterns across experimental groups. Data are shown as individual animals with mean ± SEM. n=9-12 mice/group.

### 3.4 Plasma EVs from chronic Iso/OP mice mediate pro-inflammatory responses in the hippocampus

To determine whether plasma EVs generated after Iso/OP exposure carry functional inflammatory activity, EVs isolated from Sham/F, Iso/OP/F, Sham/M, and Iso/OP/M donor mice were stereotactically injected into the hippocampus of naïve young adult male C57BL/6 recipients. Hippocampal tissue was collected 24 h after injection and analyzed using a cytokine/chemokine array to assess the acute neuroimmune response. Compared with sham-derived EVs, Iso/OP-derived EVs induced distinct hippocampal cytokine and chemokine profiles, demonstrating that chronic Iso/OP alters the inflammatory bioactivity of circulating EVs (Fig. 7A). Nine analytes exhibited clear donor sex-dependent differences between the Iso/OP/M and Iso/OP/F EV groups (Fig. 7B). Among these, PF-4/CXCL4 and IGFBP-3 showed the largest differences, while Fractalkine/CX3CL1, IL-12, GM-CSF, MIP-1α/CCL3, LIX/CXCL5, Leptin R, and MCP-5/CCL12 further indicated differential regulation of neuron-microglia signaling, myeloid activation, chemotaxis, and inflammatory responses. Together, these findings demonstrate that Iso/OP-derived plasma EVs are biologically active mediators capable of acutely reshaping the hippocampal inflammatory milieu in a donor sex-dependent manner, rather than serving solely as circulating biomarkers.

**Fig. 7.**
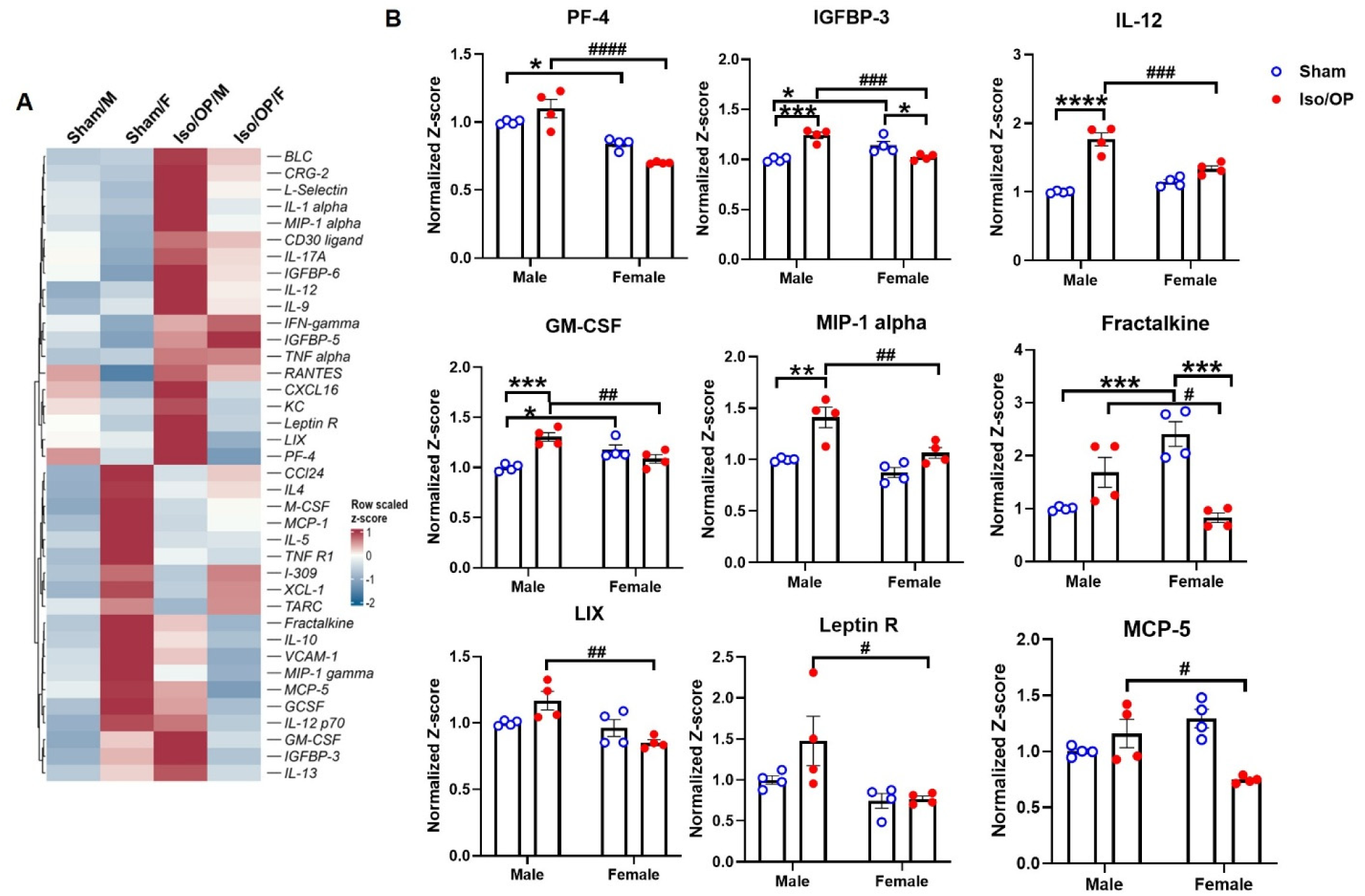
Iso/OP-derived plasma EVs induce donor sex-dependent cytokine and chemokine remodeling after hippocampal injection. Plasma EVs were isolated from male and female sham or Iso/OP mice and stereotactically injected into the hippocampus of naive young adult male C57BL/6 mice (n=5-6 mice/group). Hippocampal tissue was collected 24 h after injection and analyzed using a cytokine array membrane. **A** Heatmap showing row-scaled z-scores of hippocampal cytokine-and chemokine-related proteins after injection of EVs derived from each donor group. Proteins are clustered by abundance patterns. **B** Representative cytokine/chemokine-related proteins showing donor sex-and Iso/OP-associated differences after hippocampal EV injection. Cytokine array spot intensities were normalized to positive control spots on each membrane and then normalized to the male sham EV-injected group. Duplicate spots from two independent membranes were combined for each analyte. Heatmap and bar graph values are shown as row-scaled z-scores derived from normalized array intensities. Data are shown as individual animals with mean ± SEM. Kruskal-Wallis p values are shown for overall group comparisons; selected pairwise comparisons are indicated above plots. Two-way ANOVA p values for donor sex, Iso/OP, and sex x Iso/OP interaction are shown below each plot.

## 4 Discussion

The present study demonstrates that Iso/OP exposure induces persistent, sex-dependent neurological and systemic alterations that extend beyond the perioperative period. Twelve weeks after exposure, Iso/OP mice exhibited sustained olfactory deficits, with female mice showing additional impairments in odor sensitivity, spatial working memory, and associative learning. These behavioral changes were accompanied by sex-dependent transcriptomic remodeling in the OB and HI and persistent alterations in circulating plasma EVs, including reduced EV abundance and sex-dependent changes in EV protein and inflammatory cargo. Importantly, EVs from Iso/OP mice triggered acute hippocampal cytokine and chemokine responses in naïve recipients, demonstrating their capacity to modulate the neuroimmune environment. Together, these findings identify Iso/OP as a sustained systemic stressor that drives sex-dependent behavioral, molecular, and EV-mediated inflammatory changes.

Postoperative neurological dysfunction has primarily been examined during acute or subacute recovery periods, with most preclinical studies assessing cognitive outcomes from hours to several weeks after Iso exposure or anesthesia/surgery [59–62]. Whether extended volatile anesthetic exposure combined with surgery induces behavioral abnormalities that persist for months in young adult wild-type mice remains unclear. This distinction is important because early behavioral changes may reflect transient anesthetic effects, perioperative inflammation, pain, or stress, whereas deficits persisting to 12 weeks are more likely to indicate lasting neurobiological remodeling. Our laparotomy model provides a clinically relevant surgical inflammatory stimulus, while prolonged Iso exposure broadens the translational relevance to extended volatile anesthetic administration. Although not intended to model intensive care unit sedation directly, prolonged inhaled Iso has been used for critical care sedation through anesthetic-conserving delivery systems [8, 63]. This paradigm therefore enabled us to determine whether extended anesthesia/surgical stress produces persistent neurological deficits and whether these chronic outcomes differ by biological sex.

Previous studies have shown that volatile anesthetics and surgery can impair cognition, but they vary widely in animal age, sex, exposure duration, follow-up period, and behavioral endpoints. Most have focused on short-term outcomes, aged animals, male-only cohorts, or disease-prone models. Lin et al. reported that Iso exposure induced hippocampal injury, neuroinflammation, and cognitive impairment in adult mice, supporting the concept that volatile anesthetics can directly affect hippocampal-dependent function [59], although the study did not address long-term outcomes or sex as a biological variable. The concentration and duration of Iso exposure influenced later cognitive performance, reinforcing the importance of anesthetic exposure duration as an experimental variable [60, 61]. Aged, but not adult, mice showed spatial learning deficits after 2 h Iso exposure or appendectomy, emphasizing age as a major modifier of postoperative cognitive outcomes [64]. The absence of detectable effects of sex or Iso exposure on cognitive/behavioral performance or amyloid-related biomarkers at an early disease stage in Tg2576 mice [65] suggests that the role of sex remains incompletely understood. Our study fills this important knowledge gap by demonstrating that extended Iso/OP induces long-term neurological and behavioral deficits in young adult mice in a sex-dependent manner.

The olfactory findings build on emerging evidence linking postoperative olfactory dysfunction with cognitive impairment. Elderly patients undergoing abdominal surgery with general anesthesia showed olfactory impairment and cognitive decline on postoperative days 3 and 7, with olfactory impairment correlating with short-term and delayed memory deficits [66]. Similarly, anesthesia/surgery induced both olfactory and cognitive deficits in mice, which were alleviated by odor enrichment, supporting a functional link between olfactory dysfunction and postoperative neurocognitive impairment [67]. Sex differences in olfactory function are also well documented, with females generally exhibiting greater odor sensitivity and discrimination than males, although these effects vary by species and experimental conditions [68–71]. Extending previous studies, our findings demonstrate that extended Iso/OP induces behavioral abnormalities that persist for at least 12 weeks in young adult wild-type C57BL/6 mice. While both sexes exhibited lasting olfactory impairment, female mice also developed deficits in odor sensitivity, spatial working memory, and associative learning, suggesting that biological sex influences the long-term neurological consequences of perioperative stress.

The olfactory deficits observed after chronic Iso/OP were accompanied by long-term transcriptomic remodeling in the OB, suggesting that the behavioral phenotype is associated with durable molecular alterations in olfactory sensory circuit organization and signaling. Notably, several of the affected pathways have direct relevance to OB-dependent odor processing. MAPK/ERK signaling has been implicated in olfactory sensory neuron survival and odor-guided behavior, as odorant stimulation can activate MAPK/CREB-dependent survival pathways in olfactory sensory neurons, while conditional ERK5 deletion reduces odor-detection sensitivity and short-term odor memory [72, 73]. Although the present RNAseq analysis was performed in OB tissue rather than olfactory epithelium, Iso/OP-associated alterations in MAPK-related pathways may still reflect disruption of glomerular afferent signaling, local OB circuit adaptation, or downstream odor-processing mechanisms. In parallel, odorant receptor activation can engage MAPK-and Rho-dependent signaling pathways, supporting the relevance of the small GTPase/Rho-related signatures identified in the OB after Iso/OP [74]. These molecular changes may be particularly important because olfactory sensory axons converge onto discrete glomerular targets, and altered axon guidance, cell projection organization, or glomerular maintenance could impair odor coding and odor sensitivity [75]. In addition, the enrichment of synaptic organization and synaptic vesicle-related pathways is consistent with impaired OB circuit function, as granule-cell-mediated inhibition of mitral cells is critical for rapid and accurate discrimination of similar odors [76]. Together, these findings suggest that Iso/OP-induced olfactory dysfunction, particularly the greater impairment observed in female mice, may involve disruption of OB pathways governing synaptic communication, glomerular circuit organization, and activity-dependent sensory processing.

The HI transcriptomic findings provide a molecular context for the sex-dependent cognitive deficits observed after chronic Iso/OP. Because the hippocampus is critical for spatial working memory and associative learning, its function relies on coordinated regulation of synaptic plasticity, extracellular matrix (ECM) remodeling, metabolism, and neuroimmune signaling. Hippocampal plasticity is also sexually dimorphic, suggesting that males and females may engage distinct molecular responses to the same perioperative exposure [77]. Notably, ECM remodeling regulates dendritic spine stability and memory-related circuit plasticity [78], while chemokine signaling can directly impair hippocampal function. For example, MIP-1α/CCL3 suppresses hippocampal synaptic transmission and LTP, leading to deficits in spatial and long-term memory [79]. Thus, the broader transcriptomic changes in Iso/OP female hippocampus likely reflect coordinated disruption of synaptic, ECM, metabolic, and immune pathways that contribute to their greater cognitive vulnerability.

Iso/OP also produced chronic remodeling of circulating plasma EVs, extending our findings from brain-region-specific molecular changes to systemic extracellular signaling. This is notable because EVs were collected 12 weeks after Iso/OP, indicating that these changes do not simply represent acute perioperative debris or transient postoperative inflammation. Prior work has shown that anesthetics and surgery can alter EV composition and function [80], and surgical trauma alone can modify circulating EV cargo in mice [81]. Clinically, altered circulating EV cargo has also been associated with cognitive decline after major surgery [82]. Our own prior studies further support the concept that acute CNS insults-induced circulating EVs can carry neuroinflammatory potential [31, 55, 83], and chronic spinal cord injury produced sex-dependent EV responses associated with distal brain neuroinflammation and neurodegeneration [48]. In the perioperative context, our recent multiorgan transcriptomic and circulating EV profiling study showed that Iso/OP induces age-dependent systemic vulnerability, with circulating EV profiles mirroring tissue-level shifts and EV protein cargo enriched for inflammatory and growth factor-related signatures in older mice [26]. Consistent with this framework, the present study shows that Iso/OP reduces plasma EV concentration, alters the small EV-enriched size range, and reshapes EV-associated protein and inflammatory cargo in a sex-dependent manner at the chronic phase.

Importantly, EV characterization and cargo profiling alone cannot establish whether these persistent circulating alterations have functional consequences within the brain. To address this question, the hippocampal injection experiment directly tested whether plasma EVs collected 12 weeks after Iso/OP retained pro-inflammatory bioactivity. Hippocampal administration of Iso/OP-derived EVs into naïve recipient mice elicited distinct cytokine and chemokine responses, demonstrating that these EVs are sufficient to acutely alter the hippocampal inflammatory milieu. This observation is consistent with recent evidence showing that circulating EVs altered by anesthesia and surgery can transmit pathological signals, as administration of anesthesia/surgery-derived EVs induced delirium-like behavioral deficits and neuronal pathology in aged mice [84]. In the present study, the marked donor sex-dependent differences in PF-4/CXCL4 and IGFBP-3 suggest differential regulation of platelet/vascular-associated and growth factor-related signaling pathways. In contrast, differences in Fractalkine/CX3CL1, IL-12, GM-CSF, MIP-1α/CCL3, LIX/CXCL5, Leptin R, and MCP-5/CCL12 point to broader alterations in neuroimmune communication, myeloid activation, chemotactic signaling, and metabolic-inflammatory pathways. Notably, because all recipient mice were naïve males, the observed hippocampal cytokine responses are most likely attributable to donor sex-dependent differences in EV bioactivity rather than to recipient sex. Collectively, these findings indicate that plasma EVs generated during the chronic phase following Iso/OP are not merely persistent circulating biomarkers, but biologically active mediators capable of modulating hippocampal inflammatory signaling long after the initial perioperative insult.

In conclusion, this study identifies extended isoflurane anesthesia combined with laparotomy as a durable systemic and neurobiological stressor that induces chronic, sex-dependent neurological alterations in young adult mice. Twelve weeks after exposure, Iso/OP resulted in sustained olfactory dysfunction, female-biased impairment in hippocampal-dependent behaviors, sex-specific transcriptomic remodeling in both the OB and HI, and long-lasting changes in the abundance and inflammatory cargo of circulating plasma EVs. Notably, plasma EVs isolated during this chronic phase retained functional inflammatory bioactivity, as evidenced by their ability to acutely alter hippocampal cytokine and chemokine responses following intracerebral administration into naïve recipient mice. These findings support a model in which extended anesthesia and surgical stress establish a persistent peripheral EV signature capable of modulating central inflammatory signaling long after the initial perioperative insult. Collectively, our results highlight biological sex as a critical determinant of long-term neuroimmune responses to anesthesia and surgery and identify circulating EVs as both promising biomarkers and potential mediators of long-term postoperative neurological vulnerability.

## Abbreviations

BF: Buried food
CNS: Central nervous system
DEG: differential expression gene
EV: Extracellular vesicles
ECM: Extracellular matrix
F: Female
HI: Hippocampus
Iso: Isoflurane
M: Male
OB: Olfactory bulb
OP: Operation
OM: Odor memory
PLS-DA: partial least squares discriminant analysis RNAseq RNA sequencing
YM: Y maze

## Authors’ contributions

Concept creation and study design: JW, YL, RP; Data collection: YL, RP, BK, ZL, H. Li; Data analysis: YL, BK, ZL, H. Lee; Manuscript writing: YL, JW, RP. All authors have critically read and commented on the final paper.

## Data availability

The datasets used and/or analyzed during the current study are available from the corresponding author on reasonable request.

## Declarations

### Ethics approval and consent to participate

Animal experiments were carried out according to experimental protocols approved by the Animal Care and Use Committee (IACUC) of the University of Maryland School of Medicine.

### Consent for publication

Not applicable.

## References

1. Kim HY, Lee JE, Kim HY, Kim J: Volatile sedation in the intensive care unit: A systematic review and meta-analysis. Medicine (Baltimore) 2017, 96:e8976.

2. Breaux AM, Miller GR, Cooper HD, Bembenick KN, Reddy A, Ahmadzadeh S, Shekoohi S, Kaye AD: Critical Care Sedation: Emerging Clinical Considerations and Risks of Volatile Anesthetics for Sedation: A Narrative Review. Diseases 2026, 14.

3. Perouansky M, Hemmings HC, Jr.: Neurotoxicity of general anesthetics: cause for concern? Anesthesiology 2009, 111:1365–1371.

4. Evered L, Silbert B, Knopman DS, Scott DA, DeKosky ST, Rasmussen LS, Oh ES, Crosby G, Berger M, Eckenhoff RG, Nomenclature Consensus Working G: Recommendations for the nomenclature of cognitive change associated with anaesthesia and surgery-2018. Br J Anaesth 2018, 121:1005–1012.

5. Wang J, Liu Z: Research progress on molecular mechanisms of general anesthetic-induced neurotoxicity and cognitive impairment in the developing brain. Front Neurol 2022, 13:1065976.

6. Ries CR, Azmudeh A, Franciosi LG, Schwarz SK, MacLeod BA: Cost comparison of sevoflurane with isoflurane anesthesia in arthroscopic menisectomy surgery. Can J Anaesth 1999, 46:1008–1013.

7. Moody AE, Beutler BD, Moody CE: Predicting cost of inhalational anesthesia at low fresh gas flows: impact of a new generation carbon dioxide absorbent. Med Gas Res 2020, 10:64–66.

8. Meiser A, Volk T, Wallenborn J, Guenther U, Becher T, Bracht H, Schwarzkopf K, Knafelj R, Faltlhauser A, Thal SC, et al: Inhaled isoflurane via the anaesthetic conserving device versus propofol for sedation of invasively ventilated patients in intensive care units in Germany and Slovenia: an open-label, phase 3, randomised controlled, non-inferiority trial. Lancet Respir Med 2021, 9:1231–1240.

9. Jabaudon M, Zhai R, Blondonnet R, Bonda WLM: Inhaled sedation in the intensive care unit. Anaesth Crit Care Pain Med 2022, 41:101133.

10. Bajwa NM, Lee JB, Halavi S, Hartman RE, Obenaus A: Repeated isoflurane in adult male mice leads to acute and persistent motor decrements with long-term modifications in corpus callosum microstructural integrity. J Neurosci Res 2019, 97:332–345.

11. Miatello J, Palacios-Cuesta A, Radell P, Oberthuer A, Playfor S, Amores-Hernandez I, Barreault S, Biedermann R, Charlo Molina MT, Encarnacion Martinez J, et al: Inhaled isoflurane for sedation of mechanically ventilated children in intensive care (IsoCOMFORT): a multicentre, randomised, active-control, assessor-masked, non-inferiority phase 3 trial. Lancet Respir Med 2025, 13:897–910.

12. Lin D, Zuo Z: Isoflurane induces hippocampal cell injury and cognitive impairments in adult rats. Neuropharmacology 2011, 61:1354–1359.

13. Zuo CL, Wang CM, Liu J, Shen T, Zhou JP, Hao XR, Pan YZ, Liu HC, Lian QQ, Lin H: Isoflurane anesthesia in aged mice and effects of A1 adenosine receptors on cognitive impairment. CNS Neurosci Ther 2018, 24:212–221.

14. Sneyd JR: Avoiding kidney damage in ICU sedation with sevoflurane: use isoflurane instead. Br J Anaesth 2022, 129:7–10.

15. Gentz BA, Malan TP, Jr.: Renal toxicity with sevoflurane: a storm in a teacup? Drugs 2001, 61:2155–2162.

16. Assefi M, Chiarito A, Blanchard F, Baron E, Clavieras N, James A, Constantin JM: Renal Impact of Prolonged Sevoflurane Sedation in Intensive Care Unit Patients: An Observational Study. Am J Crit Care 2025, 34:439–448.

17. Moller JT, Cluitmans P, Rasmussen LS, Houx P, Rasmussen H, Canet J, Rabbitt P, Jolles J, Larsen K, Hanning CD, et al: Long-term postoperative cognitive dysfunction in the elderly ISPOCD1 study. ISPOCD investigators. International Study of Post-Operative Cognitive Dysfunction. Lancet 1998, 351:857–861.

18. Zhao S, Wang B, Liu M, Yu D, Li J: The impact of preoperative frailty on perioperative neurocognitive disorders in elderly patients: A systematic review and meta-analysis. J Res Med Sci 2024, 29:47.

19. Nakagoshi N, Locatelli FM, Kitamura S, Hirota S, Kawano T: The impact of preoperative stress on age-related cognitive dysfunction after abdominal surgery: a study using a rat model. BMC Res Notes 2024, 17:369.

20. Peng W, Lu W, Jiang X, Xiong C, Chai H, Cai L, Lan Z: Current Progress on Neuroinflammation-mediated Postoperative Cognitive Dysfunction: An Update. Curr Mol Med 2023, 23:1077–1086.

21. Li Y, Uzun C, Islam ST, Krishnamachary B, Lee H, Wang Z, Li H, Liu S, Wu J: Age-dependent Transcriptional and Circuit Alterations in the brain Underlie Post-Anesthesia Neurobehavioral Dysfunction. Aging Dis 2025.

22. Fodale V, Tripodi VF, Penna O, Fama F, Squadrito F, Mondello E, David A: An update on anesthetics and impact on the brain. Expert Opin Drug Saf 2017, 16:997–1008.

23. Colon E, Bittner EA, Kussman B, McCann ME, Soriano S, Borsook D: Anesthesia, brain changes, and behavior: Insights from neural systems biology. Prog Neurobiol 2017, 153:121–160.

24. Avidan MS, Evers AS: Review of clinical evidence for persistent cognitive decline or incident dementia attributable to surgery or general anesthesia. J Alzheimers Dis 2011, 24:201–216.

25. Maniaci A, Lentini M, Trombadore R, Gruppuso L, Milardi S, Scrofani R, Cuttone G, Sorbello M, Modica R, Lechien JR, et al: Neurological and Olfactory Disturbances After General Anesthesia. Life (Basel) 2025, 15.

26. Krishnamachary B, Lee H, Wang Z, Yang WW, Li H, Lei Z, Wu J, Li Y: Multiorgan transcriptomics and circulating extracellular vesicle profiling reveal age-dependent systemic vulnerability to isoflurane anesthesia and surgery. Geroscience 2026.

27. Thery C, Witwer KW, Aikawa E, Alcaraz MJ, Anderson JD, Andriantsitohaina R, Antoniou A, Arab T, Archer F, Atkin-Smith GK, et al: Minimal information for studies of extracellular vesicles 2018 (MISEV2018): a position statement of the International Society for Extracellular Vesicles and update of the MISEV2014 guidelines. J Extracell Vesicles 2018, 7:1535750.

28. Hansen DR, Svenningsen P: New guidelines to uncover the physiology of extracellular vesicles. Acta Physiol (Oxf) 2024, 240:e14153.

29. Cabrera-Pastor A: Extracellular Vesicles as Mediators of Neuroinflammation in Intercellular and Inter-Organ Crosstalk. Int J Mol Sci 2024, 25.

30. Kumar MA, Baba SK, Sadida HQ, Marzooqi SA, Jerobin J, Altemani FH, Algehainy N, Alanazi MA, Abou-Samra AB, Kumar R, et al: Extracellular vesicles as tools and targets in therapy for diseases. Signal Transduct Target Ther 2024, 9:27.

31. Khan NZ, Cao T, He J, Ritzel RM, Li Y, Henry RJ, Colson C, Stoica BA, Faden AI, Wu J: Spinal cord injury alters microRNA and CD81+ exosome levels in plasma extracellular nanoparticles with neuroinflammatory potential. Brain Behav Immun 2021, 92:165–183.

32. Dutta D, Khan N, Wu J, Jay SM: Extracellular Vesicles as an Emerging Frontier in Spinal Cord Injury Pathobiology and Therapy. Trends Neurosci 2021, 44:492–506.

33. Wei FS, Rao MW, Huang YL, Chen SB, Wu YQ, Yang L: miR-182-5p Delivered by Plasma Exosomes Promotes Sevoflurane-Induced Neuroinflammation and Cognitive Dysfunction in Aged Rats with Postoperative Cognitive Dysfunction by Targeting Brain-Derived Neurotrophic Factor and Activating NF-kappaB Pathway. Neurotox Res 2022, 40:1902–1912.

34. Piao L, Na OH, Seo EH, Hong SW, Sohn KM, Kwon Y, Lee SH, Kim SH: Effects of general anaesthesia with an inhalational anaesthetic agent on the expression of exosomes in rats. Int J Med Sci 2022, 19:1399–1407.

35. Valadi H, Ekstrom K, Bossios A, Sjostrand M, Lee JJ, Lotvall JO: Exosome-mediated transfer of mRNAs and microRNAs is a novel mechanism of genetic exchange between cells. Nat Cell Biol 2007, 9:654–659.

36. Wang H, Guo X, Zhu X, Li Y, Jia Y, Zhang Z, Yuan S, Yan F: Gender Differences and Postoperative Delirium in Adult Patients Undergoing Cardiac Valve Surgery. Front Cardiovasc Med 2021, 8:751421.

37. Oh ES, Sieber FE, Leoutsakos JM, Inouye SK, Lee HB: Sex Differences in Hip Fracture Surgery: Preoperative Risk Factors for Delirium and Postoperative Outcomes. J Am Geriatr Soc 2016, 64:1616–1621.

38. Wiredu K, Mueller A, McKay TB, Behera A, Shaefi S, Akeju O: Sex Differences in the Incidence of Postoperative Delirium after Cardiac Surgery: A Pooled Analyses of Clinical Trials. Anesthesiology 2023, 139:540–542.

39. Zhang C, Zhang Y, Shen Y, Zhao G, Xie Z, Dong Y: Anesthesia/Surgery Induces Cognitive Impairment in Female Alzheimer’s Disease Transgenic Mice. J Alzheimers Dis 2017, 57:505–518.

40. Lee BH, Chan JT, Kraeva E, Peterson K, Sall JW: Isoflurane exposure in newborn rats induces long-term cognitive dysfunction in males but not females. Neuropharmacology 2014, 83:9–17.

41. Tuohy KP, Tynan AR, Gerisma JN, Marx JO: Effect of Mouse (Mus musculus) Sex and C57BL/6 Substrain on Sensitivity to Isoflurane and Ketamine-Xylazine-Acepromazine Anesthesia. J Am Assoc Lab Anim Sci 2025, 64:1–10.

42. Yang WW, Chen AW, Lee H, Li H, Lee JG, Li Y, Shen WB: Isoflurane and Surgical Stress Disrupt Fatty Acid and Carbon Metabolism, Leading to Cardiomyopathy in Aged Mice. Cells 2026, 15.

43. Tillerson JL, Caudle WM, Parent JM, Gong C, Schallert T, Miller GW: Olfactory discrimination deficits in mice lacking the dopamine transporter or the D2 dopamine receptor. Behav Brain Res 2006, 172:97–105.

44. Liu X, Lei Z, Gilhooly D, He J, Li Y, Ritzel RM, Li H, Wu LJ, Liu S, Wu J: Traumatic brain injury-induced inflammatory changes in the olfactory bulb disrupt neuronal networks leading to olfactory dysfunction. Brain Behav Immun 2023, 114:22–45.

45. Yang WW, Matyas JJ, Li Y, Lee H, Lei Z, Renn CL, Faden AI, Dorsey SG, Wu J: Dissecting Genetic Mechanisms of Differential Locomotion, Depression, and Allodynia after Spinal Cord Injury in Three Mouse Strains. Cells 2024, 13.

46. Ritzel RM, Li Y, Jiao Y, Doran SJ, Khan N, Henry RJ, Brunner K, Loane DJ, Faden AI, Szeto GL, Wu J: Bi-directional neuro-immune dysfunction after chronic experimental brain injury. J Neuroinflammation 2024, 21:83.

47. Ritzel RM, Li Y, Jiao Y, Lei Z, Doran SJ, He J, Shahror RA, Henry RJ, Khan R, Tan C, et al: Brain injury accelerates the onset of a reversible age-related microglial phenotype associated with inflammatory neurodegeneration. Sci Adv 2023, 9:eadd1101.

48. Li Y, Khan N, Ritzel RM, Lei Z, Allen S, Faden AI, Wu J: Sexually dimorphic extracellular vesicle responses after chronic spinal cord injury are associated with neuroinflammation and neurodegeneration in the aged brain. J Neuroinflammation 2023, 20:197.

49. Li Y, Ritzel RM, Lei Z, Cao T, He J, Faden AI, Wu J: Sexual dimorphism in neurological function after SCI is associated with disrupted neuroinflammation in both injured spinal cord and brain. Brain Behav Immun 2022, 101:1–22.

50. Macheda T, Snider HC, Watson JB, Roberts KN, Bachstetter AD: An active avoidance behavioral paradigm for use in a mild closed head model of traumatic brain injury in mice. J Neurosci Methods 2020, 343:108831.

51. Dobin A, Davis CA, Schlesinger F, Drenkow J, Zaleski C, Jha S, Batut P, Chaisson M, Gingeras TR: STAR: ultrafast universal RNA-seq aligner. Bioinformatics 2013, 29:15– 21.

52. Li B, Dewey CN: RSEM: accurate transcript quantification from RNA-Seq data with or without a reference genome. BMC Bioinformatics 2011, 12:323.

53. Love MI, Huber W, Anders S: Moderated estimation of fold change and dispersion for RNA-seq data with DESeq2. Genome Biol 2014, 15:550.

54. Wu T, Hu E, Xu S, Chen M, Guo P, Dai Z, Feng T, Zhou L, Tang W, Zhan L, et al: clusterProfiler 4.0: A universal enrichment tool for interpreting omics data. Innovation (Camb) 2021, 2:100141.

55. Lei Z, Krishnamachary B, Khan NZ, Ji Y, Li Y, Li H, Brunner K, Faden AI, Jones JW, Wu J: Spinal cord injury disrupts plasma extracellular vesicles cargoes leading to neuroinflammation in the brain and neurological dysfunction in aged male mice. Brain Behav Immun 2024.

56. Huang Y, Arab T, Russell AE, Mallick ER, Nagaraj R, Gizzie E, Redding-Ochoa J, Troncoso JC, Pletnikova O, Turchinovich A, et al: Toward a human brain extracellular vesicle atlas: Characteristics of extracellular vesicles from different brain regions, including small RNA and protein profiles. Interdiscip Med 2023, 1:e20230016.

57. Ritchie ME, Phipson B, Wu D, Hu Y, Law CW, Shi W, Smyth GK: limma powers differential expression analyses for RNA-sequencing and microarray studies. Nucleic Acids Res 2015, 43:e47.

58. Smyth GK: Linear models and empirical bayes methods for assessing differential expression in microarray experiments. Stat Appl Genet Mol Biol 2004, 3:Article3.

59. Lin D, Cao L, Wang Z, Li J, Washington JM, Zuo Z: Lidocaine attenuates cognitive impairment after isoflurane anesthesia in old rats. Behav Brain Res 2012, 228:319– 327.

60. Liu J, Wang P, Zhang X, Zhang W, Gu G: Effects of different concentration and duration time of isoflurane on acute and long-term neurocognitive function of young adult C57BL/6 mouse. Int J Clin Exp Pathol 2014, 7:5828–5836.

61. Miao H, Dong Y, Zhang Y, Zheng H, Shen Y, Crosby G, Culley DJ, Marcantonio ER, Xie Z: Anesthetic Isoflurane or Desflurane Plus Surgery Differently Affects Cognitive Function in Alzheimer’s Disease Transgenic Mice. Mol Neurobiol 2018, 55:5623–5638.

62. Valentim AM, Alves HC, Olsson IA, Antunes LM: The effects of depth of isoflurane anesthesia on the performance of mice in a simple spatial learning task. J Am Assoc Lab Anim Sci 2008, 47:16–19.

63. Sackey PV, Martling CR, Granath F, Radell PJ: Prolonged isoflurane sedation of intensive care unit patients with the Anesthetic Conserving Device. Crit Care Med 2004, 32:2241–2246.

64. Foteinou PT, Greenstein JL, Winslow RL: Mechanistic Investigation of the Arrhythmogenic Role of Oxidized CaMKII in the Heart. Biophys J 2015, 109:838–849.

65. Su H, Bo Y, Zhang X, Zhang J, Gao Z, Yu Z: Associations of folate intake with all-cause and cause-specific mortality among individuals with diabetes. Front Nutr 2022, 9:1021709.

66. Lan Y, You ZJ, Du R, Chen LS, Wu JX: Association of Olfactory Impairment and Postoperative Cognitive Dysfunction in Elderly Patients. Front Mol Biosci 2021, 8:681463.

67. Zhang C, Han Y, Liu X, Tan H, Dong Y, Zhang Y, Liang F, Zheng H, Crosby G, Culley DJ, et al: Odor Enrichment Attenuates the Anesthesia/Surgery-induced Cognitive Impairment. Ann Surg 2023, 277:e1387–e1396.

68. Lillqvist M, Claeson AS, Zakrzewska M, Andersson L: Comparable responses to a wide range of olfactory stimulation in women and men. Sci Rep 2023, 13:9059.

69. Bontempi C, Jacquot L, Brand G: Sex Differences in Odor Hedonic Perception: An Overview. Front Neurosci 2021, 15:764520.

70. Sorokowski P, Karwowski M, Misiak M, Marczak MK, Dziekan M, Hummel T, Sorokowska A: Sex Differences in Human Olfaction: A Meta-Analysis. Front Psychol 2019, 10:242.

71. Doty RL, Cameron EL: Sex differences and reproductive hormone influences on human odor perception. Physiol Behav 2009, 97:213–228.

72. Watt WC, Sakano H, Lee ZY, Reusch JE, Trinh K, Storm DR: Odorant stimulation enhances survival of olfactory sensory neurons via MAPK and CREB. Neuron 2004, 41:955–967.

73. Pan YW, Kuo CT, Storm DR, Xia Z: Inducible and targeted deletion of the ERK5 MAP kinase in adult neurogenic regions impairs adult neurogenesis in the olfactory bulb and several forms of olfactory behavior. PLoS One 2012, 7:e49622.

74. Benbernou N, Esnault S, Galibert F: Activation of SRE and AP1 by olfactory receptors via the MAPK and Rho dependent pathways. Cell Signal 2013, 25:1486–1497.

75. Treloar HB, Feinstein P, Mombaerts P, Greer CA: Specificity of glomerular targeting by olfactory sensory axons. J Neurosci 2002, 22:2469–2477.

76. Abraham NM, Egger V, Shimshek DR, Renden R, Fukunaga I, Sprengel R, Seeburg PH, Klugmann M, Margrie TW, Schaefer AT, Kuner T: Synaptic inhibition in the olfactory bulb accelerates odor discrimination in mice. Neuron 2010, 65:399–411.

77. Wang W, Le AA, Hou B, Lauterborn JC, Cox CD, Levin ER, Lynch G, Gall CM: Memory-Related Synaptic Plasticity Is Sexually Dimorphic in Rodent Hippocampus. J Neurosci 2018, 38:7935–7951.

78. Ferrer-Ferrer M, Dityatev A: Shaping Synapses by the Neural Extracellular Matrix. Front Neuroanat 2018, 12:40.

79. Marciniak E, Faivre E, Dutar P, Alves Pires C, Demeyer D, Caillierez R, Laloux C, Buee L, Blum D, Humez S: The Chemokine MIP-1alpha/CCL3 impairs mouse hippocampal synaptic transmission, plasticity and memory. Sci Rep 2015, 5:15862.

80. Abel F, Giebel B, Frey UH: Agony of choice: How anesthetics affect the composition and function of extracellular vesicles. Adv Drug Deliv Rev 2021, 175:113813.

81. Mkrtchian S, Ebberyd A, Veerman RE, Mendez-Lago M, Gabrielsson S, Eriksson LI, Gomez-Galan M: Surgical Trauma in Mice Modifies the Content of Circulating Extracellular Vesicles. Front Immunol 2021, 12:824696.

82. Mkrtchian S, Eldh M, Ebberyd A, Gabrielsson S, Vegvari A, Ricksten SE, Danielson M, Oras J, Wiklund A, Eriksson LI, Gomez-Galan M: Changes in circulating extracellular vesicle cargo are associated with cognitive decline after major surgery: an observational case-control study. Br J Anaesth 2025, 134:1683–1695.

83. Park C, Lei Z, Li Y, Ren B, He J, Huang H, Chen F, Li H, Brunner K, Zhu J, et al: Extracellular vesicles in sepsis plasma mediate neuronal inflammation in the brain through miRNAs and innate immune signaling. J Neuroinflammation 2024, 21:252.

84. Gao Y, Tang S, Liu J, Yang X, Chen H, Li J, Ni X: The Administration of Circulating Extracellular Vesicles Modified by Anesthesia and Surgery Induces Delirium-Like Behaviors in Aged Mice. CNS Neurosci Ther 2025, 31:e70483.

