## Supplemental Materials for "Sex-dependent chronic neurological dysfunction following isoflurane anesthesia and surgery is associated with circulating extracellular vesicle-mediated neuroinflammatory signaling"

#Ms. Ruth Park is a third-year medical student at University of Maryland School of Medicine, Baltimore, MD 21201 USA

**Supplementary Information**

Supplemental Information includes three Supplemental figures and figure legends.

**Supplemental Fig. S1**

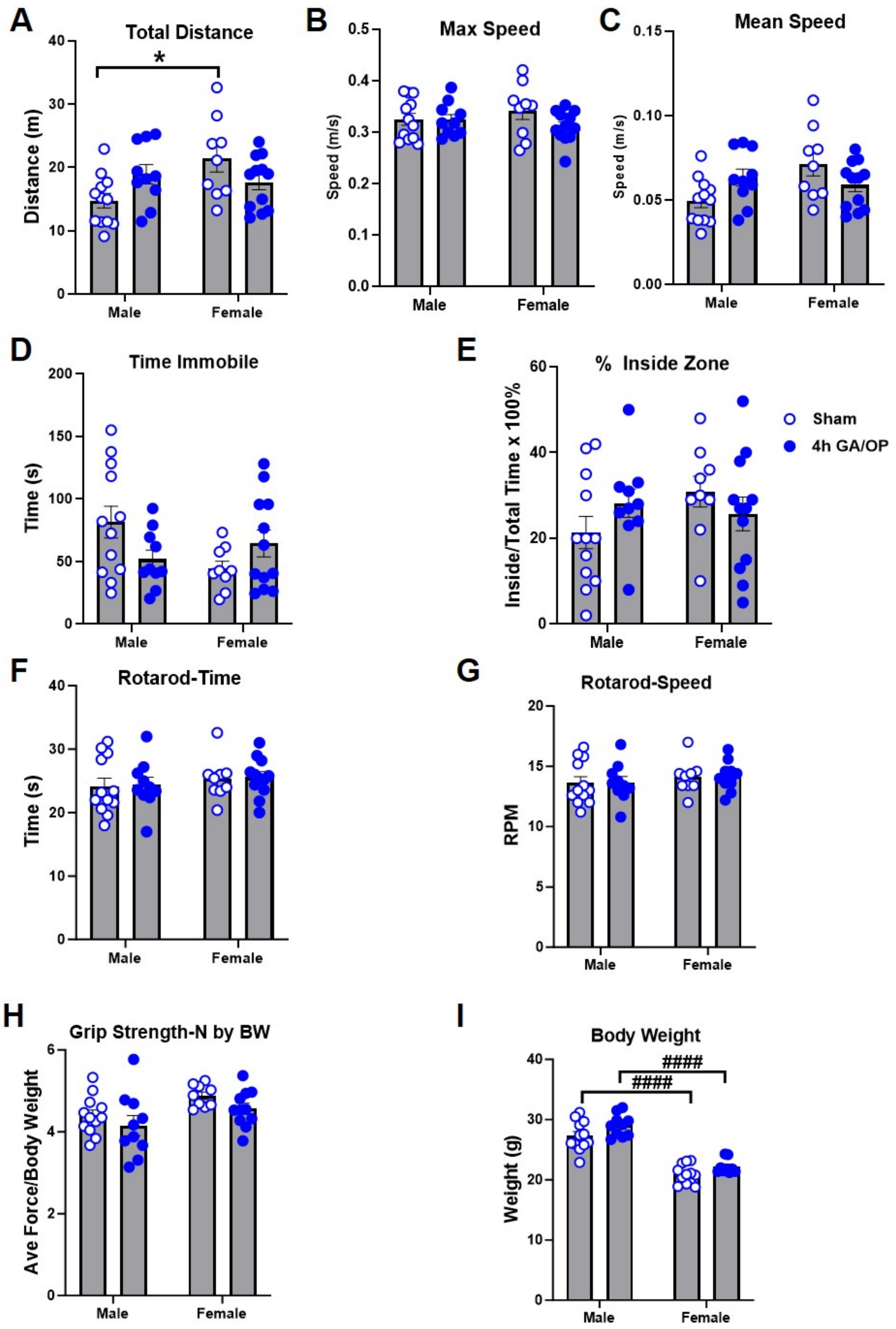

**Supplemental Fig. S1. Locomotor activity, motor function, grip strength, and body weight following 4 h isoflurane and surgery.** Male and female mice were assessed at 3 months following Sham or 4 h isoflurane and surgery (Iso/OP). **A-E** Open-field activity was evaluated by total distance travelled **(A)**, maximum speed **(B)**, mean speed **(C)**, time spent immobile **(D)**, and percentage of total time spent within the inside zone **(E)**. **F-G** Motor coordination was assessed by rotarod test, as recorded by latency to fall **(F)** from the accelerating rotarod and rotarod speed at fall **(G)**. **(H)** Grip strength was normalized to body weight. **(I)** Body weight of male and female mice. Open circles indicate Sham mice and filled circles indicate 4 h Iso/OP mice. Individual points represent individual animals, and bars represent mean ± SEM. *$p < 0.05$; #### $p < 0.0001$ for the indicated comparisons. Data were analyzed by two-way ANOVA with Tukey's multiple-comparisons test. n = 9-12 mice/group.

**Supplemental Fig. S2**

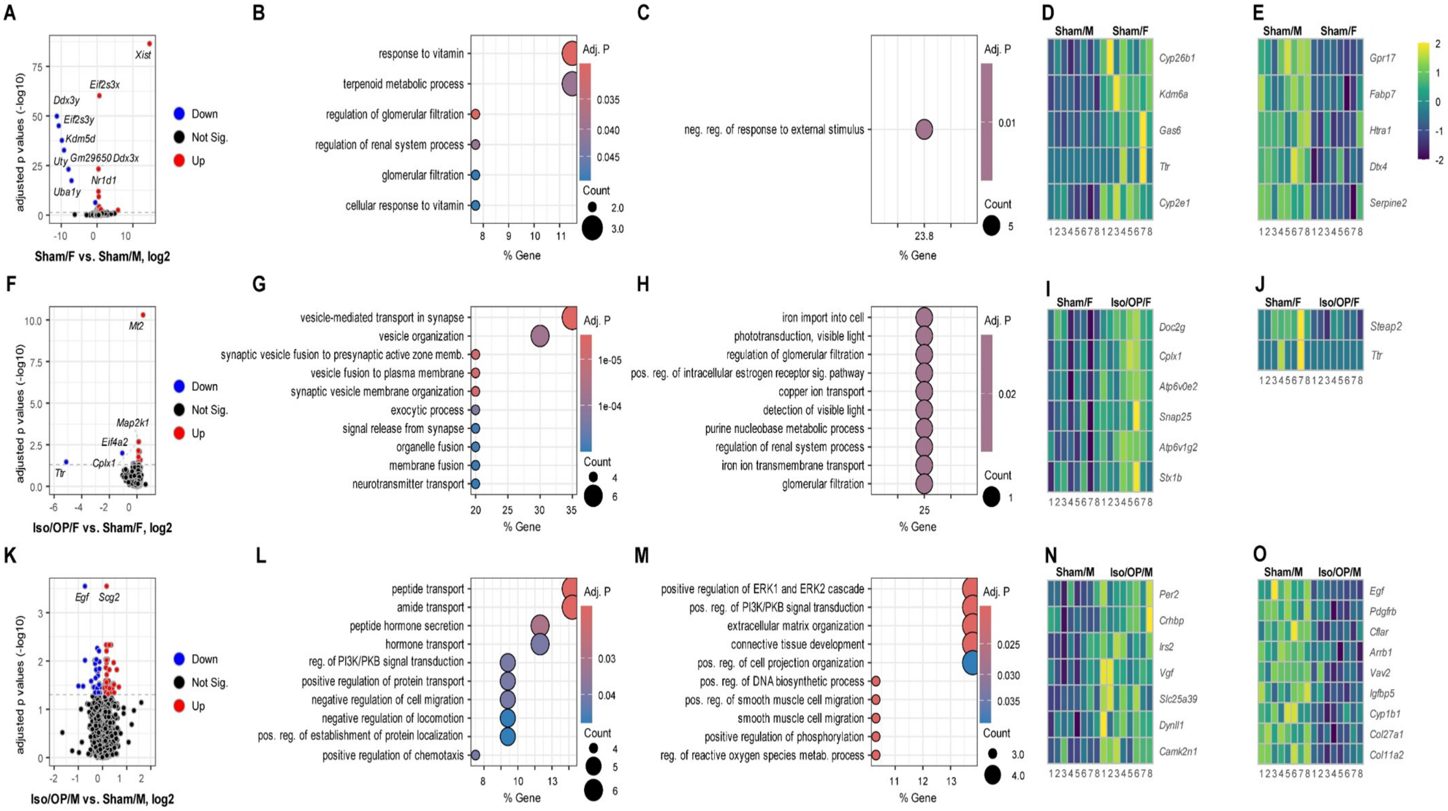


**Supplemental Fig. S2. Baseline sex-associated differences and within-sex Iso/OP responses in OB transcriptomics at 3 months after exposure. A-E** Baseline sex-associated differences in sham mice. Volcano plot showing DEGs in Sham/Female vs. Sham/Male mice **(A)**, GO biological process enrichment of upregulated and downregulated DEGs **(B-C)**, and heatmap visualization of representative genes from selected enriched pathways **(D-E)**. **F-J** Within-female Iso/OP response. Volcano plot showing DEGs in Iso/OP/Female vs. Sham/Female mice **(F)**, GO biological process enrichment of upregulated and downregulated DEGs **(G-H)**, and heatmap visualization of representative genes from selected enriched pathways **(I-J)**. **K-O** Within-male Iso/OP response. Volcano plot showing DEGs in Iso/OP/Male vs. Sham/Male mice **(K)**, GO biological process enrichment of upregulated and downregulated DEGs **(L-M)**, and heatmap visualization of representative genes from selected enriched pathways **(N-O)**. Dot size represents gene count, color indicates adjusted p value, and the x-axis shows gene ratio. RNA-seq data were analyzed with DESeq2; n=8 mice/group.

### Supplemental Fig. S3

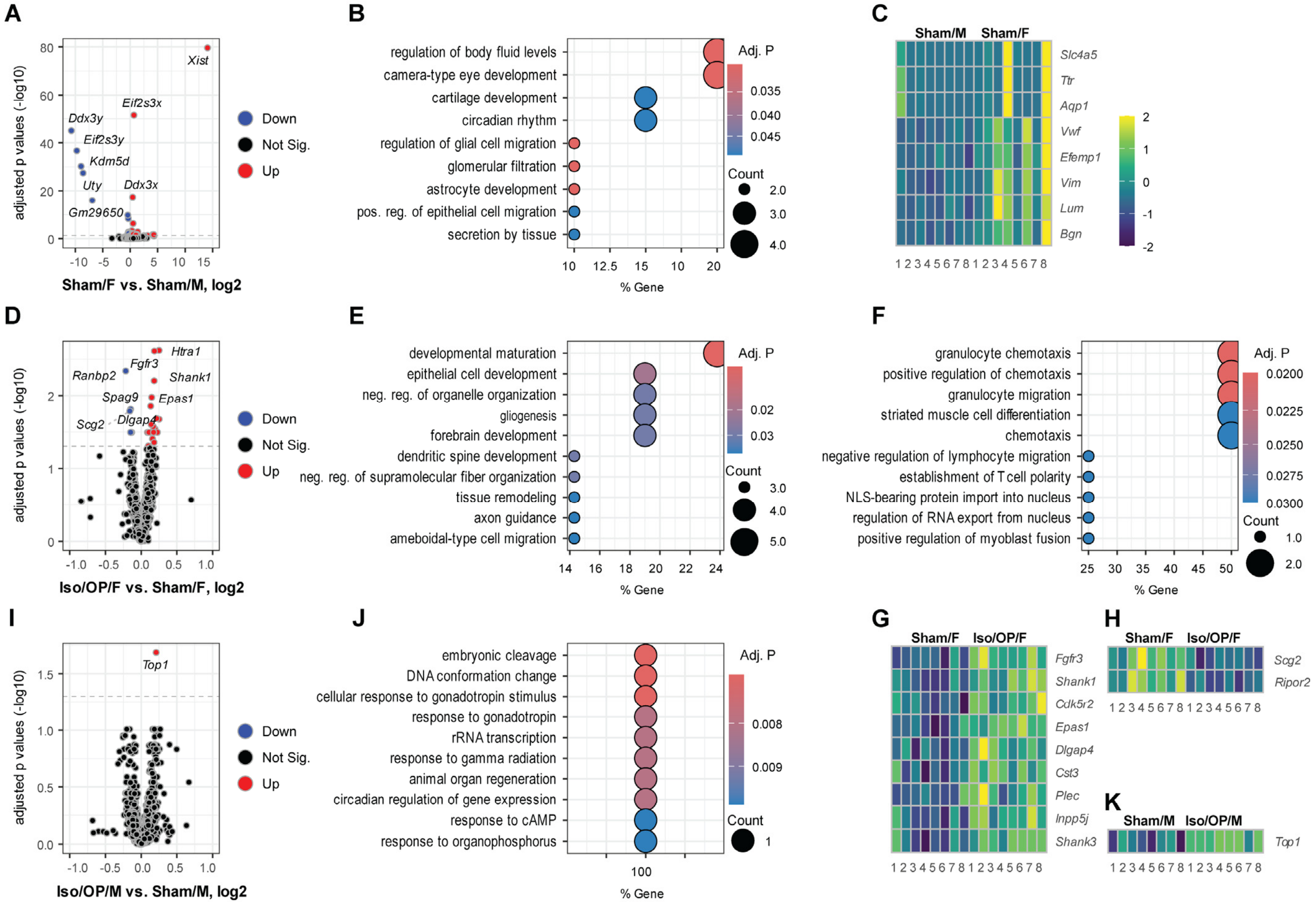


**Supplemental Fig. S3. Baseline sex-associated differences and within-sex Iso/OP responses in HI transcriptomics at 3 months after exposure**. **A-C** Baseline sex-associated differences in sham mice. Volcano plot showing DEGs in Sham/Female vs. Sham/Male mice **(A)**, GO biological process enrichment of baseline sex-associated DEGs **(B)**, and heatmap visualization of representative transcripts **(C)**. **D-H** Within-female Iso/OP response. Volcano plot showing DEGs in Iso/OP/Female vs. Sham/Female mice **(D)**, GO biological process enrichment of upregulated and downregulated DEGs **(E-F)**, and heatmap visualization of representative upregulated and downregulated genes **(G-H)**. **I-K** Within-male Iso/OP response. Volcano plot showing DEGs in Iso/OP/Male vs. Sham/Male mice **(I)**, GO biological process enrichment/annotation of male Iso/OP-associated DEGs **(J)**, and heatmap visualization of *Top1* expression across Sham/Male and Iso/OP/Male samples **(K)**. Dot size represents gene count, color indicates adjusted p value, and the x-axis shows gene ratio. RNA-seq data were analyzed with DESeq2; n=8 mice/group.
